# Structural Basis for HIV-1 Maturation Inhibition by PF-46396 Determined by MAS NMR

**DOI:** 10.1101/2025.07.15.664917

**Authors:** Roman Zadorozhnyi, Caitlin M. Quinn, Kaneil K. Zadrozny, Sherimay D. Ablan, Brandon J. Kennedy, Glenn P. A. Yap, Daniel Sanner, Christina Kraml, Eric O. Freed, Barbie K. Ganser-Pornillos, Owen Pornillos, Angela M. Gronenborn, Tatyana Polenova

**Affiliations:** Department of Chemistry and Biochemistry, University of Delaware, Newark, DE 19716, United States; Pittsburgh Center for HIV Protein Interactions, University of Pittsburgh School of Medicine, 1051 Biomedical Science Tower 3, 3501 Fifth Avenue, Pittsburgh PA 15261, United States; Department of Molecular Physiology and Biological Physics, University of Virginia School of Medicine, Charlottesville, VA 22908, United States; HIV Dynamics and Replication Program, Center for Cancer Research, National Cancer Institute, Frederick, MD 21702, United States; Lotus Separations LLC, Princeton University, Princeton, NJ 08540, United States; Department of Biochemistry, University of Utah, Salt Lake City, UT 84112, United States; Department of Structural Biology, University of Pittsburgh School of Medicine, Pittsburgh, PA 15261 USA

**Keywords:** HIV-1, maturation inhibitors, PF-46396, inositol hexakisphosphate, magic angle spinning (MAS) NMR, atomic-resolution structure

## Abstract

Among the different types of HIV-1 maturation inhibitors, those that stabilize the junction between the capsid protein C-terminal domain (CA_CTD_) and the spacer peptide 1 (SP1) within the immature Gag lattice are promising candidates for antiretroviral therapies. Here, we report the atomic-resolution structure of CA_CTD_-SP1 assemblies with the small-molecule maturation inhibitor PF- 46396 and the assembly cofactor inositol hexakisphosphate (IP6), determined by magic angle spinning (MAS) NMR spectroscopy. Our results reveal that although the two PF-46396 enantiomers exhibit distinct binding modes, they both possess similar anti-HIV potency. PF-46396 binding arrests IP6 dynamics in the six-helix bundle pore, and the two enantiomers induce unique IP6 orientations in the pore. Importantly, our data suggest the presence of monoanionic IP6 form IP6 in the complex. Our study establishes the structural basis for PF-46396 action and suggests a mechanistic model for drug resistance.

## INTRODUCTION

The Gag polyprotein in the immature HIV-1 virion comprises the linked matrix (MA), capsid (CA), nucleocapsid (NC), and p6 domains, along with two spacer peptides, SP1 and SP2^1–3^. During HIV-1 maturation, the viral protease cleaves the Gag polyprotein into its constituent subdomains in a stepwise manner^4^, resulting in a structural reorganization within the viral particle. In the final step of this process, SP1 is cleaved from the CA-SP1 intermediate, initiating the generation of mature cores of infectious virions. Maturation inhibitors (MIs) are anti-retroviral compounds that block virus maturation^5–11^. MIs have been categorized into three classes: molecules that (i) interfere with CA-SP1 processing, (ii) prevent capsid condensation, and (iii) promote integrase aggregation and interfere with viral particle formation in late replication (ALLINIs)^12^.

In immature HIV-1 virions, the Gag polyprotein forms a hexagonal lattice via its CA domain^13^, with residues 225-231 of CA’s C-terminal domain (CA_CTD_), together with SP1, folding into a six-helix bundle (6HB)^14, 15^ (Fig. 1). Efficient Gag lattice assembly requires inositol hexakisphosphate (IP6), a key cofactor stabilizing the 6HB^16–18^. The cleavage site for the viral protease (PR) is located at the CA_CTD_-SP1 junction and requires a partial unfolding of the 6HB. The cleavage is required to form mature conical capsids. MIs that interfere with CA-SP1 cleavage stabilize the 6HB and thereby reduce the processing rate. The first-in-class MI, Bevirimat (BVM), a betulinic acid derivative, exhibits this mode of action^6, 7^, and the atomic-level details of its interactions with the immature Gag lattice and CA_CTD_-SP1 assemblies have been recently elucidated^15, 19, 20^. Unfortunately, certain subtypes of HIV-1 that exhibit amino acid sequence polymorphisms in CA and SP1 are poorly inhibited by BVM, rendering this compound unsuitable for antiretroviral therapy in individuals living with HIV^21–23^. However, BVM has been valuable as a scaffold in the development of second-generation analogs, which exhibit higher potency and broader antiviral activity against HIV compared to BVM^24–28^.

**Figure 1.**
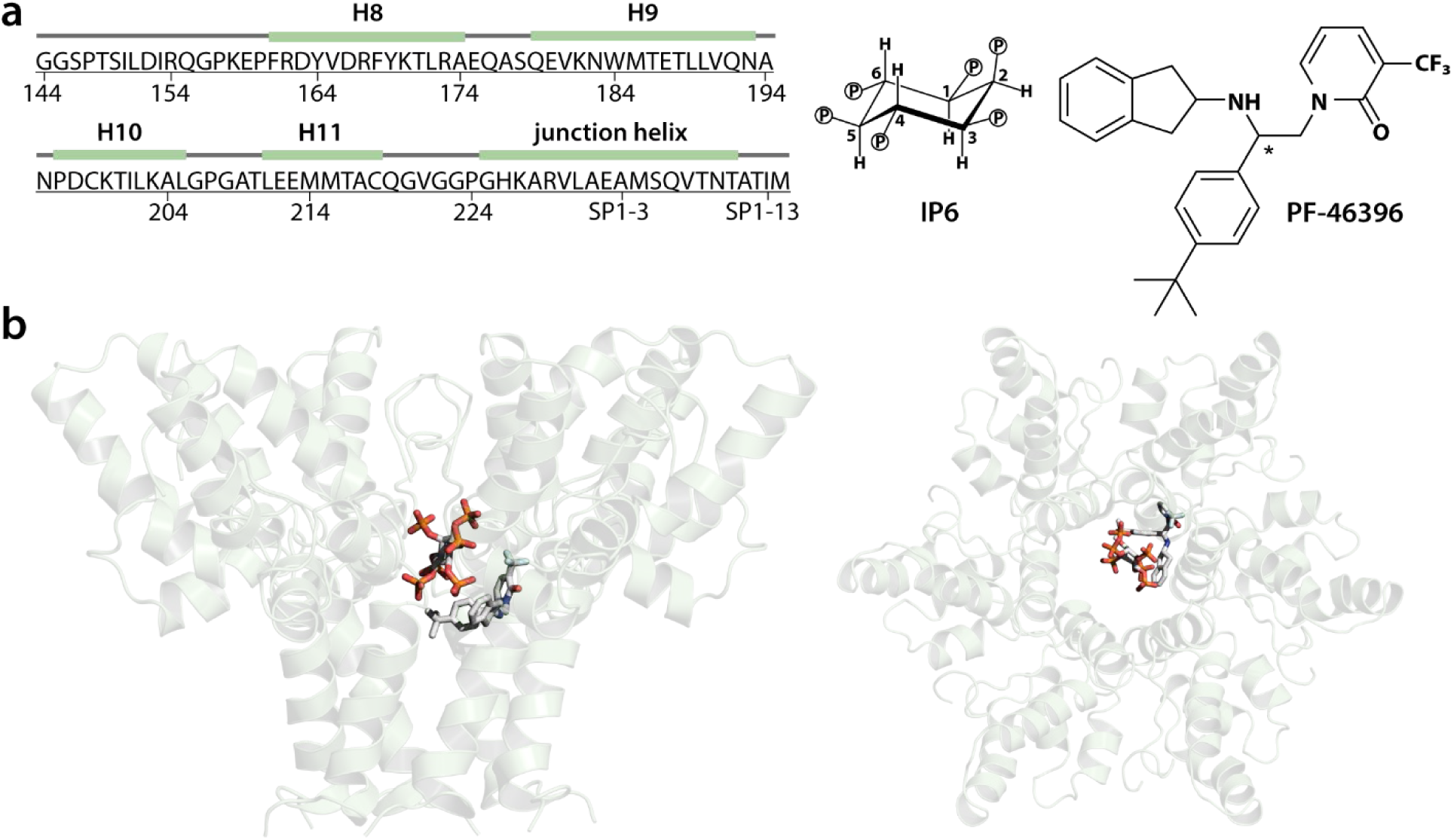
| CA_CTD_-SP1 structure in the presence of PF-46396. **a)** Amino acid sequence of CA_CTD_-SP1 with secondary structure elements marked (left) and chemical structures of IP6 and PF-46396 molecules (right). The chiral carbon atom in PF-46396 is labeled (*). **b)** Cartoon representation of the MAS NMR structure of a single hexamer of CA_CTD_-SP1 in the presence of PF-46396(*R*) and IP6 in stick representation: side view (left) and top view (right).

A different MI with an alternate scaffold is 1-[2-(4-tert-butylphenyl)-2-(2,3-dihydro-1H-inden- 2-ylamino)ethyl]-3-(trifluoromethyl)pyridine-2-one or PF-46396^29^. PF-46396 is a propeller-shaped molecule with three distinct types of chemical moieties (Fig. 1a). The structure of PF-46396 and activity towards BVM-resistant HIV-1 variants, like SP1-V7A^30^, suggested that an alternative mode of binding to the immature lattice may be responsible for this enhanced activity. We, therefore, pursued structural studies on the CA_CTD_-SP1/IP6 complex with PF-46396.

Here, we report the atomic-resolution structure of crystalline CA_CTD_-SP1 assemblies in the presence of PF-46396 and IP6, determined by magic angle spinning nuclear magnetic resonance (MAS NMR) spectroscopy. We characterized the binding modes of racemic PF-46396 and pure PF-46396*(S)* and PF-46396*(R)* enantiomers. Our results reveal that PF-46396 binds asymmetrically inside the CA_CTD_-SP1 hexamer’s 6HB, with unique binding poses for the two enantiomers. Remarkably, binding of the individual PF-46396 enantiomers also results in distinct orientations of IP6 in the pore of the 6HB. Our structural models suggest a mechanistic explanation for PF-46396-induced drug resistance and may guide the future design of more potent maturation inhibitors.

## EXPERIMENTAL

### Protein Expression and Purification

The expression plasmid for the CA_CTD_-SP1 fragment of HIV-1 strain NL4-3 Gag, containing a non-cleavable His-tag and encoding the P373T change (SP1-P10T) was described previously^14^. Proteins were expressed in transformed *E. coli* BL21 (DE3) cells, which were grown at 37 °C until mid-log phase (OD_600_ of 0.6-1), induced with 1 mM IPTG and further grown overnight at 18 °C. Expression of U-^13^C,^15^N enriched and fractionally deuterated (FD) U-^13^C,^15^N enriched CA_CTD_-SP1 proteins comprised additional pre-culturing steps to slowly adapt the cells from rich medium to minimal medium, as reported previously^35, 36^. Cells were harvested by centrifugation and stored at −80 °C until use.

Protein purification was performed as described previously^14^. In brief, bacterial pellets were resuspended in 50 mM Tris, pH 8.3, 1 M LiCl, containing protease inhibitor tablets (Roche) and supplemented with 0.3% (w/v) deoxycholate. Cells were lysed by incubation with lysozyme, followed by sonication. Lysates were clarified by centrifugation, filtered, and incubated with Ni-NTA agarose resin (Qiagen) for 30 min at 4 °C. Unbound fractions were washed away, and bound protein was eluted using a step gradient of 15 mM to 300 mM imidazole. Protein was purified to homogeneity using anion exchange chromatography in 20 mM Tris, pH 8.0, 0.5 M NaCl. Pure protein was concentrated to 10 mg/mL, flash-frozen in liquid nitrogen, and stored at −80 °C until use.

### Buffer Exchange

The FD-^13^C,^15^N-CA_CTD_-SP1 sample was prepared in 20 mM Tris, pD 8.0, 0.5 M NaCl buffer made from 1 M Tris stock buffer in 99.9% D_2_O (Cambridge Isotope Laboratories, Inc) with the pD adjusted with deuterium chloride (Sigma-Aldrich) in 99.9% D_2_O (Cambridge Isotope Laboratories, Inc). In this FD protocol, the only source of deuterium is D_2_O, while the other components of the growth media are protonated. Most of the exchangeable protons in the resulting protein samples are deuterated. Final protein samples were prepared by buffer exchange as follows: 0.5 mL protein at 10 mg/mL was diluted into 10 mL with 20 mM Tris, pD 8.0, 0.5 M NaCl buffer and concentrated using the centrifugal filter units, 4 times. After exchange, the samples were recovered in a final volume 0.5 mL of 20 mM Tris, pD 8.0, 0.5 M NaCl buffer, with protein concentrations ranging from 9 to 10 mg/mL.

### Protein Assembly and Sample Preparation

Proteins were assembled at 0.4-0.5 mM with equimolar IP6 (Sigma-Aldrich) and PF-46396 (Aobious, Inc) in a final volume of 1-2 mL. Assemblies were incubated overnight at 16-20 °C. Assemblies (approximately 50 mg protein) were centrifuged at 10000 x g and packed in 3.2 mm thin-wall and 1.9 mm Bruker rotors. 1.3 mm Bruker rotors were packed at 20000 x g using an ultracentrifuge rotor packing tool.

### PF-46396 Chiral Separation

A chiral supercritical fluid chromatography method was developed by Lotus Separations, LLC (Princeton, NJ). Preparative separation of PF-46396 enantiomers was achieved using a Berger Multi-gram II SFC instrument equipped with two Varian SD-1 pumps, a Knauer K-2501 spectrophotometer, and a 6-ton bulk CO_2_ tank with a built-in chiller, heater, and G700 compressor. Optimized preparative chromatography conditions were obtained on a ColumnTek EnantioCel^®^ C2-5 SFC column (21 x 250 mm; 5 µm ColumnTek, LLC, State College, PA). Scaled preparative conditions were performed at 70 mL/min flowrate with a mobile phase of 35% methanol, 65% CO_2_ (bone-dry grade), a column temperature of 35 °C, and the column backpressure set to 100 bar.

### PF-46396 Antiviral Activity and Particle Infectivity

The activity of PF-46396 against the SP1-P10 and SP1-T10 viral derivatives was determined essentially as reported previously^37^. Briefly, HEK293T cells (ATCC, Cat# CRL-3216) were transfected with pNL4-3/SP1-P10 or pNL4-3/SP1-T10 molecular clones, and the transfected cells were treated with 24 concentrations of PF-46396 ranging from 0 to 10 μM. Virus-containing supernatants were harvested, normalized for reverse transcriptase (RT) activity, and used to infect the TZM-bl indicator cell line. Infectivity data from three independent experiments were analyzed with GraphPad Prism 7. Curves were fit using nonlinear regression as log (inhibitor) versus normalized response, with a variable slope using a least-squares (ordinary) fit.

### MAS NMR Spectroscopy

MAS NMR experiments on U-^13^C,^15^N-CA_CTD_-SP1/PF-46396(*racem*)/IP6, U-^13^C,^15^N-CA_CTD_- SP1/PF-46396(*R*)/IP6, and U-^13^C,^15^N-CA_CTD_-SP1/PF-46396(*S*)/IP6 were performed at 14.1 T on a Bruker AVIII HD spectrometer equipped with a standard-bore Magnex magnet, outfitted with a 3.2 mm E-Free HCN probe. The Larmor frequencies were 599.8 MHz (^1^H), 150.8 MHz (^13^C), and 60.8 MHz (^15^N). The actual sample temperature was calibrated using KBr as an external temperature sensor (by recording the ^79^Br T_1_ relaxation time)^38^ and was maintained at 4±1 °C using the Bruker temperature controller. The MAS frequency was 14 kHz, controlled to within ±10 Hz by a Bruker MAS controller. The typical 90° pulse lengths were 2.4 μs (^1^H), 2.9 μs (^13^C), and 4.6 μs (^15^N). For ^1^H-^13^C and ^1^H-^15^N cross-polarization (CP), a linear 20% amplitude ramp was applied on ^1^H, with the center of the ramp matched to the Hartmann-Hahn condition at the first spinning sideband; the ^1^H-^13^C and ^1^H-^15^N contact times were 1.2 ms and 1.4 ms, respectively. The typical radio frequency (rf) fields during ^1^H-^13^C and ^1^H-^15^N CP were 66 kHz (^1^H), 49 kHz (^13^C), and 50 kHz (^15^N). The 2D combined R2_n_^ν^-driven (CORD)^39^ spectra were recorded with mixing times of 50 ms, 100 ms, 250 ms, 500 ms, with judiciously chosen sets for different samples; the ^1^H field strength during CORD was 14 kHz. The contact time for the band-selective ^15^N-^13^C magnetization transfer was 6.5 ms. The rf fields for NCA transfer were 35 kHz (^13^C) and 21 kHz (^15^N). SPINAL-64^40^ decoupling (80 kHz) was used during the evolution (t_1_) and acquisition (t_2_) times. A 2D phase-shifted ^13^C-detected proton-assisted insensitive-nuclei cross polarization (PAIN-CP)^41^ experiment was also acquired on the U-^13^C,^15^N-CA_CTD_-SP1/PF-46396(*racem*)/IP6 sample. During the PAIN-CP mixing period, rf field strengths for ^1^H, ^13^C and ^15^N channels were all 60 kHz. The PAIN-CP mixing time was 4.6 ms.

^1^H-detected MAS NMR experiments (hNH-dREDOR and ^1^H-^1^H-RFDR-dREDOR) on the U- ^13^C,^15^N-CA_CTD_-SP1/PF-46396(*racem*)/IP6 sample were performed at 14.1 T on a Bruker AVIII HD spectrometer equipped with a standard-bore Magnex magnet and outfitted with a 1.3 mm HCN probe. The MAS frequency was 60 kHz controlled to within ±10 Hz by a Bruker MAS controller. The actual sample temperature was calibrated with KBr as described above and maintained at 33±1 °C throughout the experiments. The 90° pulse lengths were 1.4 μs for ^1^H and 2.8 μs for ^13^C, and 3.5 μs for ^15^N. In the hNH-dREDOR experiment, ^1^H-^15^N cross-polarization employed a linear 20% amplitude ramp on ^1^H, with the center of the ramp matched to the Hartmann-Hahn condition at the first spinning sideband; the contact time was 10 ms. 10 kHz WALTZ-16^42^ ^13^C and ^15^N decoupling was applied during the ^1^H acquisition period. 15 kHz SWf-TPPM^43^ ^1^H decoupling was applied during the ^15^N evolution period. MISSISSIPPI^44^ water suppression was applied before the dephasing period. MAS NMR double-REDOR filtered hNH-dREDOR and ^1^H-^1^H-RFDR-dREDOR experiments employed simultaneous ^1^H-^13^C/^1^H-^15^N REDOR^45^ dephasing periods of 3 and 10 ms, respectively, to eliminate signals from protons directly bound to ^13^C and ^15^N. Radiofrequency-driven recoupling (RFDR^46^) utilized 5.56 μs π-pulses; the mixing time was 10 ms.

^1^H-detected experiments on FD-^13^C,^15^N-CA_CTD_-SP1/PF-46396(*racem*)/IP6, as well as ^31^P-detected MAS NMR experiments on U-^13^C,^15^N-CA_CTD_-SP1/PF-46396(*racem*)/IP6, CA_CTD_-SP1/PF- 46396(*R*)/IP6 and CA_CTD_-SP1/PF-46396(*S*)/IP6 crystalline arrays were performed at 20.0 T, on a Bruker AVIII spectrometer equipped with a standard-bore Bruker Ascend magnet and outfitted with a 1.9 mm HX probe. The typical 90° pulse lengths were 2.1 μs for ^1^H, 3.1 μs for ^13^C and 3.5 μs for ^31^P. The ^1^H-^13^C and ^1^H-^31^P cross polarization employed a linear 20% amplitude ramp on ^1^H, and the center of the ramp was matched to the Hartmann-Hahn condition at the first spinning sideband, with contact times of 10 and 3.5 ms, respectively. The MAS frequency was 14 or 40 kHz, controlled to within ±10 Hz by a Bruker MAS controller. The actual sample temperature was calibrated with KBr as described above and maintained at 15±1 °C for the experiments conducted at 40 kHz MAS, and -10±1 °C for the experiments performed at 14 kHz MAS, using the Bruker temperature controller.

^19^F- and ^13^C-detected MAS NMR experiments on the U-^13^C,^15^N-CA_CTD_-SP1/PF- 46396(*racem*)/IP6 sample were performed at 11.7 T on a Bruker AVANCE III spectrometer outfitted with a wide-bore Magnex magnet, equipped with a 1.3 mm HFX probe. The Larmor frequencies were 500.1 MHz for ^1^H, 470.6 MHz for ^19^F, and 125.8 MHz for ^13^C. The MAS frequency was 60 kHz controlled to within ±10 Hz by a Bruker MAS controller. The actual sample temperature was calibrated with KBr as described above and maintained at 37 ± 1 °C throughout the experiments. The typical 90° pulse lengths were 2.1 μs for ^1^H, 2.5 μs for ^19^F, and 3.2 μs for ^13^C. The ^1^H-^19^F cross-polarization employed a linear 20% amplitude ramp on ^1^H, with the center of the ramp matched to the Hartmann-Hahn condition at the first spinning sideband; the contact times were 2.5 ms and 10 ms. The transferred echo double resonance (TEDOR^47^) mixing time was 1.5 ms.

### Solution NMR Spectroscopy

Europium (III) tris[3-(heptafluoropropylhydroxymethilene)-*d*-camphorate (Eu(hfc)_3_) was purchased from Sigma-Aldrich. ^1^H and ^13^C solution NMR spectra of PF-46396(*racem*)/DMSO-D6, PF-46396(*racem*)/Eu(hfc)_3_/DMSO-D6 and IP6/D2O were collected at 14.1 T (^1^H Larmor frequency of 600.1 MHz) on a Bruker Avance spectrometer equipped with a triple-resonance inverse detection (TXI) probe.

### Data Processing

All MAS NMR spectra were processed using Bruker TopSpin and NMRPipe^48^. The ^13^C and ^15^N signals were referenced with respect to the external standards adamantane and ammonium chloride, respectively. ^1^H was indirectly referenced to the ^13^C of adamantane^49^. ^31^P was referenced with respect to the phosphorous resonance of 85% H_3_PO_4_ at 0 ppm. ^19^F was referenced to trifluoroacetic acid (100 μM solution in 25 mM NaPi, pH 6.5) used as an external reference (0 ppm). The 2D data sets were processed by applying 30°, 45°, 60° and 90° shifted sine bell apodization, followed by a Lorentzian-to-Gaussian transformation in both dimensions. Forward linear prediction to twice the number of the original data points was used in the indirect dimension in some data sets, followed by zero filling.

### Diffraction Data Collection

PF-46396 crystallization was carried out by slow evaporation of a saturated solution in hexane. A single crystal was mounted using viscous oil onto a plastic mesh and cooled to 100 K using nitrogen gas. Data were collected on a D8 Venture Photon diffractometer with Cu-Kα radiation (λ = 1.54178 Å) focused with Goebel mirrors. Unit cell parameters were obtained from fast scan data frames, 1°/s ω, of an Ewald hemisphere. The unit-cell dimensions, equivalent reflections and systematic absences in the diffraction data are uniquely consistent with *P*2_1_/*n*. The data were treated with multi-scan absorption corrections^50^. Structures were solved using intrinsic phasing methods^51^ and refined with full-matrix, least-squares procedures on *F*^2^.^52^

Non-hydrogen atoms were refined with anisotropic displacement parameters. The amine H-atom was initially located from the difference map and allowed to refine to position with *U_iso_* constrained to 1.2 N *U_eq_*. All other hydrogen atoms were treated as idealized contributions with geometrically calculated positions and with *U_iso_* equal to 1.2 *U_eq_* (1.5 *U_eq_* for methyl) of the attached atom. Atomic scattering factors are contained in the SHELXTL program library^50^.

PF-46396 (Fig. S5, Supporting Information) has the 2-pyridone moiety interacting, via the carbonyl oxygen, with the pendant ethylamine by forming H-bonded chains, propagated by a 2_1_ screw parallel to the b-axis (Fig. S6, Supporting Information). A search on the Cambridge Structural Database (ver. 5.42)^53^ for compounds with a N-aminoethyl-pyrid-2-one substructure yielded nine structures total, but only three structures display chain formation by intermolecular H-bonding *via* the amine N*H* and pyridone *O*: 1-(2-ammonioethyl)-3,5-dicyano-6-oxo-4-(pyridin- 1-ium-4-yl)-1,6-dihydropyridin-2-olate tetrafluoroborate dihydrate^54^, 1’-(2-ammonioethyl)-3’,5’- dicyano-6’-oxo-1’,6’-dihydro[3,4’-bipyridin-3-ium]-2’-olate chloride dihydrate^55^, and 1’-(2- ammonioethyl)-3’,5’-dicyano-6’-oxo-1’,6’-dihydro[3,4’-bipyridin-3-ium]-2’-olate bromide^55^. In all three structures, the N-aminoethyl moiety is charged, existing as an ammonium, exhibiting shorter H-bonds, 2.939, 2.786, and 2.887 Å, respectively, compared to PF-46396, which exhibits hydrogen bonds of 3.097 Å. The centroid-centroid distances of the face-centered stacked phenyl rings of the indane (electron-rich) and pyridone-CF_3_ (electron-poor) moieties is 3.664 Å, consistent with π-π interactions^56–58^ that can further stabilize the chains (Fig. S7, Supporting Information). The mean distance of inversion related, neighboring indane rings from different chains is 3.637 Å, which is within the range for parallel offset π-π interactions. A summary of data collection and refinement details is provided in Table S2, Supporting Information. The structure has been deposited in the Cambridge Structural Database under CCDC 2421271.

### MAS NMR Chemical Shift and Distance Restraints Assignment

All spectra were analyzed using CCPN^59^ and NMRFAM-Sparky^60, 61^. Chemical shift assignments (intra-residue/sequential assignments) for the CA_CTD_SP1/IP6 crystalline arrays were previously reported^20^, and used as initial guesses for CA_CTD_-SP1/PF-46396(*racem*)/IP6 assignments using 2D CORD and NCACX MAS NMR data sets. The superposition of U-^13^C,^15^N- CA_CTD_-SP1/PF-46396(*racem*)/IP6, U-^13^C,^15^N-CA_CTD_-SP1/PF-46396(*R*)/IP6, and U-^13^C,^15^N-CA_CTD_-SP1/PF-46396(*S*)/IP6 2D CORD spectra at mixing times of 50 and 500 ms, is shown in Fig. S1 and S2, Supporting Information. *De novo* assignments of inter-residue ^13^C-^13^C, ^15^N-^13^C correlations for U-^13^C,^15^N-CA_CTD_-SP1/PF-46396(*racem*)/IP6 were performed using 2D CORD spectra (100, 250, and 500 ms mixing times), 2D NCACX (50 ms mixing time) and PAIN-CP spectra. The ^13^C-^1^H inter-residue correlations were assigned using ^1^H-detected CH HETCOR spectra.

The correlations from PF-46396 (^1^H and ^19^F) and IP6 (^1^H and ^31^P) to protein resonances were assigned from 2D HC CP HETCOR, (H)NH dREDOR-HETCOR, 1D FC TEDOR and 2D HP HETCOR spectra of FD-^13^C,^15^N-CA_CTD_-SP1/PF-46396(*racem*)/IP6, U-^13^C,^15^N-CA_CTD_-SP1/PF-46396(*racem*)/IP6, and FD-^13^C,^15^N-CA_CTD_-SP1/IP6. PF-46396 enantiomer specific IP6-protein correlations were assigned using 2D HP HETCOR spectra of U-^13^C,^15^N-CA_CTD_-SP1/PF- 46396(*R*)/IP6 and CA_CTD_-SP1/PF-46396(*S*)/IP6 assemblies. All samples and MAS NMR experimental parameters are summarized in Table S1, Supporting Information.

### Structure Calculation

*Distance restraints*: Distance restraints were obtained from the assigned cross-peaks in MAS NMR spectra. Both unambiguous and up to 4-fold ambiguous restraints were considered. Protein-protein restraints were ^13^C-^13^C and ^15^N-^13^C restraints. IP6 and PF-46396 restraints were ^1^H (PF-46396)-^13^C (protein), ^1^H (PF-46396)-^15^N (protein), ^19^F (PF-46396)-^13^C (protein), ^31^P (IP6)-^1^H (protein), ^19^F (PF-46396)-^1^H (IP6), and ^1^H (PF-46396)-^1^H (IP6) restraints.

The number of distance restraints is summarized in Table 1. Distance boundaries were initially set to 1.5-6.5 Å (4.0±2.5 Å) and 2.0-7.2 Å (4.6± 2.6Å) for intra- and inter-residue restraints, respectively, as done previously^62^. PF-46396-protein distance restraints were initially set to 2.0-7.2 Å (4.6±2.6 Å). IP6-protein and IP6-PF-46396 distance restraints were initially set as follows: 2.0-9.2 Å (5.6±3.6 Å) for ^31^P-^1^H, and 2.0-11.2 Å (6.6±4.6 Å) for ^19^F-^1^H and ^1^H-^1^H distance restraints. The phi (φ) and psi (ψ) dihedral restraints were predicted from TALOS-N^63^ using the experimental ^13^C and ^15^N chemical shifts.

**Table 1.** | Summary of MAS NMR restraints and structure statistics

| PF-46396-Protein restraints |  | <sup>1</sup>H(PF-46396)-<sup>13</sup>C(protein) | <sup>1</sup>H(PF-46396)-<sup>15</sup>N(protein) | <sup>19</sup>F(PF-46396)-<sup>13</sup>C(protein) |
| --- | --- | --- | --- | --- |
| Unambiguous |  | 8 | 1 | 10 |
| Ambiguous |  | 0 | 2 | 20 |
| IP6-Protein restraints | <sup>1</sup>H(IP6)-<sup>13</sup>C(protein) |  | <sup>31</sup>P(IP6)-<sup>1</sup>H(protein) |  |
|  | w PF-46396(R) | w PF-46396(S) | w PF-46396(R) | w PF-46396(S) |
| Unambiguous | 1 | 1 | 17 | 20 |
| IP6-PF-46396 restraints | <sup>1</sup>H(IP6)-<sup>1</sup>H(PF-46396) | <sup>1</sup>H(IP6)-<sup>19</sup>F(PF-46396) | <sup>31</sup>P(IP6)-<sup>1</sup>H(PF-46396) |  |
|  |  |  | w PF-46396(R) | w PF-46396(S) |
| Unambiguous | 4 | 1 | 2 | 0 |
| Ambiguous | 1 | 0 | 12 | 0 |
| Protein distance restraints |  | <sup>13</sup>C-<sup>13</sup>C |  | <sup>15</sup>N-<sup>13</sup>C |
| Unambiguous |  | 1662 |  | 444 |
| Intra-residue |  | 496 |  | 326 |
| Sequential (\|i-j\|=1) |  | 193 |  | 90 |
| Medium range (1<\|i-j\|≤4) |  | 492 |  | 21 |
| Long range (\|i-j\|≤5) |  | 458 |  | 7 |
| Long range (\|i-j\|≤5) (inter-chain) |  | 14 |  | 4 |
| Long range (\|i-j\|≤5) (inter-hexamer) |  | 9 |  | 0 |
| Ambiguous |  | 86 |  | 4 |
| Intra-residue |  | 22 |  | 1 |
| Sequential (\|i-j\|=1) |  | 12 |  | 1 |
| Medium range (1<\|i-j\|≤4) |  | 23 |  | 2 |
| Long range (\|i-j\|≤5) |  | 25 |  | 0 |
| Total number of unambiguous restraints |  | 2160 |  |  |
| Restrains/residue |  | 21 |  |  |
| Percent completeness |  | 41% (C-C only) |  |  |
| Dihedral angle restraints |  |  |  |  |
| φ |  | 90 |  |  |
| ψ |  | 90 |  |  |
| Structure statistics* |  |  |  |  |
| Violations (mean ± s.d.) | CA<sub>CTD-SP1/PF-46396(R)/IP6</sub> | CA<sub>CTD-SP1/PF-46396(R)/IP6inv</sub> | CA<sub>CTD-SP1/PF-46396(S)/IP6</sub> | CA<sub>CTD-SP1/PF-46396(S)/IP6inv</sub> |
| Distance restraints ≥ 7.2 Å (Å) | 0.246±0.020 | 0.253±0.023 | 0.139±0.006 | 0.144±0.006 |
| Dihedral restraints ≥ 5 ° (°) | 7.814±0.300 | 8.084±0.379 | 8.372±0.068 | 8.432±0.089 |
| Max. protein-protein distance restraint violation (Å) | 1.380 | 1.380 | 0.660 | 1.160 |
| Max. protein-ligand distance restraint violation (Å) | 1.840 | 1.830 | 1.470 | 1.520 |
| Deviations from idealized geometry |  |  |  |  |
| Bond length (Å) | 0.014±0.001 | 0.014±0.000 | 0.013±0.000 | 0.013±0.000 |
| **Bond angles (°)** | 1.503±0.062 | 1.494±0.055 | 1.380±0.012 | 1.373±0.021 |
| **Impropers (°)** | 1.687±0.378 | 1.641±0.223 | 1.423±0.013 | 1.419±0.042 |
| **Average pairwise r.m.s.d. (Å)** |  |  |  |  |
| **Heavy** | 1.2±0.2 | 1.4±0.2 | 1.0±0.1 | 0.9±0.1 |
| **Backbone (N, Cα, C)** | 1.0±0.2 | 1.0±0.1 | 0.8±0.1 | 0.8±0.1 |

\*Calculated for 10 lowest energy central hexamers.

*Refinement of the seven hexamer units with PF-46396(R)/(S) and IP6*: Single-chain structure calculation of CA_CTD_-SP1, its refinement and docking into the cryo-EM envelope were performed as described previously^20^ using Xplor-NIH version 2.53^64–66^ and UCSF Chimera^67^, respectively. After docking, the precise location and orientation of the IP6 and PF-46396 enantiomers were identified by supplementing IP6 and PF-46396 distances as well as additional distance restraints between CA_CTD_-SP1 chains and hexamer units. The calculation was seeded from single-chain CA_CTD_-SP1 coordinates calculated from the experimental MAS NMR restraints, together with the coordinates, topology, and parameters of PF-46396(*R*)/(*S*) and IP6 generated using *genLigand.py* script in the Xplor-NIH based on the small molecule crystal structures. A random initial orientation of the molecules inside a single hexamer was used. The coordinates were expanded from a single hexamer to a hexamer-of-hexamers unit containing seven hexamers (42 chains) using the symexp command in PyMol^68^.

100 structures underwent torsion angle dynamics with an annealing schedule and a final gradient minimization in Cartesian space. The PF-46396 and IP6 molecules were free to move as rigid bodies during dynamics and final minimization. Two identical simulated annealing runs starting at 3000 K were performed for 10 ps, with a time step of 1 fs. The initial velocities were randomized to achieve a Maxwell distribution at a starting temperature of 3000 K. The temperature was subsequently reduced to 25 K in steps of 25 K. At each temperature step, dynamics was run for 400 fs, with an initial time step of 1 fs.

Standard terms for bond lengths, bond angles, and impropers were used to enforce correct covalent geometry.

A cross-correlation probability distribution potential, often utilized for experimental cryo-EM density^69^, enforced/conceded the overall shape and boundary of the hexamer of hexamers with the 8 Å density map used earlier for docking. The potential was restricted to backbone atoms (N, C, CA, and O) to ensure the density boundary would not influence side chain conformations.

A torsion angle database potential^70^ and hydrogen-bond database term^71^ were employed. Approximate non-crystallographic symmetry was imposed using Xplor-NIH’s PosDiffPot term, allowing the subunits of the hexamer to differ by up to 0.5 Å.

Force constants for distance restraints were ramped from 2 to 30 kcal mol^−1^ Å^−2^. The dihedral restraint force constants were set to 10 kcal mol^−1^ rad^−2^ for high-temperature dynamics at 3000 K and 200 kcal mol^−1^ rad^−2^ during cooling. The force constants of the cross-correlation probability distribution potential were set to 50 kcal mol^−1^ during high-temperature dynamics and cooling.

After the high-temperature dynamics and cooling in dihedral space, the annealed structures were minimized using a Powell energy minimization scheme in Cartesian space. The final bundles comprised 10 lowest-energy structures of the 100 calculated ones for each of the IP6 orientations observed in the complex with PF-46396(*R*) or (*S*) enantiomers.

RMSD values were calculated using routines in the Xplor-NIH (version 2.53)^64–66^. Secondary structure elements were classified according to TALOS-N.

## RESULTS

### Structure of CA_CTD_-SP1/PF-46396 Assemblies

We have calculated the structure of the CA_CTD_-SP1 hexamer of hexamers (HOH), with IP6 and PF-46396. The central hexamer structure is illustrated in Fig. 1b. These assemblies recapitulate the immature hexagonal Gag lattice, similar to the CA_CTD_-SP1 assemblies, in the presence of BVM^20^. The structures are well-defined with 2160 protein-protein distance restraints (21 restraints per residue or 41% completeness of C-C restraints), which were unambiguously assigned from MAS NMR spectra.

A total of 11 2D MAS NMR experiments were conducted on CA_CTD_-SP1/IP6 assemblies bound with racemic PF-46396 or with each enantiomer (summarized in Table S1, Supporting Information). The spectra exhibit exceptionally high sensitivity and resolution. The protein-protein correlations detected in the spectra were found to be identical for the CA_CTD_-SP1/IP6 assemblies bound with racemic and enantiomerically pure PF-46396, but distinct from those in the assemblies lacking the MI (Fig. S1 and S2, Supporting Information). As anticipated, there are no significant structural changes upon PF-46396 binding to the assembly. The SP1 residues up to I13 are present in the spectra (Fig. S3, Supporting Information) and show no significant chemical shift perturbations (CSPs) in comparison to CA_CTD_-SP1/IP6. This result, along with the chemical shift values for the observed residues, indicates that the SP1 region remains ordered and helical in the presence of PF-46396.

2D ^13^C-^13^C MAS NMR correlation spectra for the assemblies bound with PF-46396(*racem*), PF-46396(*R*), or PF-46396(*S*) show no significant spectral differences (Fig. S1, Supporting Information). The antiviral activities of the enantiomers and the racemic PF-46396 did not differ significantly based on single-cycle infectivity assays (Fig. S4, Supporting Information).

### PF-46396 Binds Asymmetrically Inside the CA_CTD_-SP1 Six-Helix-Bundle

CA_CTD_-SP1/IP6 assemblies in the presence of PF-46396 exhibit multiple changes in the 2D CORD spectra compared to CA_CTD_-SP1/IP6 assemblies alone or in the presence of BVM^20^. These include chemical shift changes and peak splittings (Fig. 2a and Fig. S1, Supporting Information). Chemical shift perturbations are observed for resonances associated with residues in the vicinity of the IP6 binding site, P157, K158, E159, H226, K227, R229, as well as P196, C198, and V221, and suggest that the PF-46396 binding site is formed by helices 10, 11 and the junction helix in the 6HB. The observed chemical shift changes are significant, up to 0.6 ppm for ^13^C, and characteristic of side chain reorientations, but not large enough to indicate any changes in secondary structure. Notably, no chemical shift changes for SP1 residues are observed, suggesting that PF-46396 binds above the CTD-SP1 junction in the 6HB. Interestingly, upon PF- 46396 binding, many of the individual cross peaks in the spectra of CA_CTD_-SP1/IP6 assemblies are split into multiple peaks, suggesting that the 6-fold symmetry of the 6HB is broken. New peaks are associated with residues P157, K158, E159, P196, V221, H226, and R229 (Fig. 2a) in two or three of 6 chains in the 6HB. The same peak splitting is observed in CA_CTD_-SP1/IP6 assemblies in the presence of the two pure PF-46396 enantiomers, as discussed below. These new peaks suggest that PF-46396 binds asymmetrically, rather than in the center of the 6HB, and interacts with a subset (two or three) of the CA_CTD_-SP1 chains. In contrast, CA_CTD_-SP1 resonances of residues that are not in contact with PF-46396 do not exhibit any chemical shift changes.

**Figure 2.**
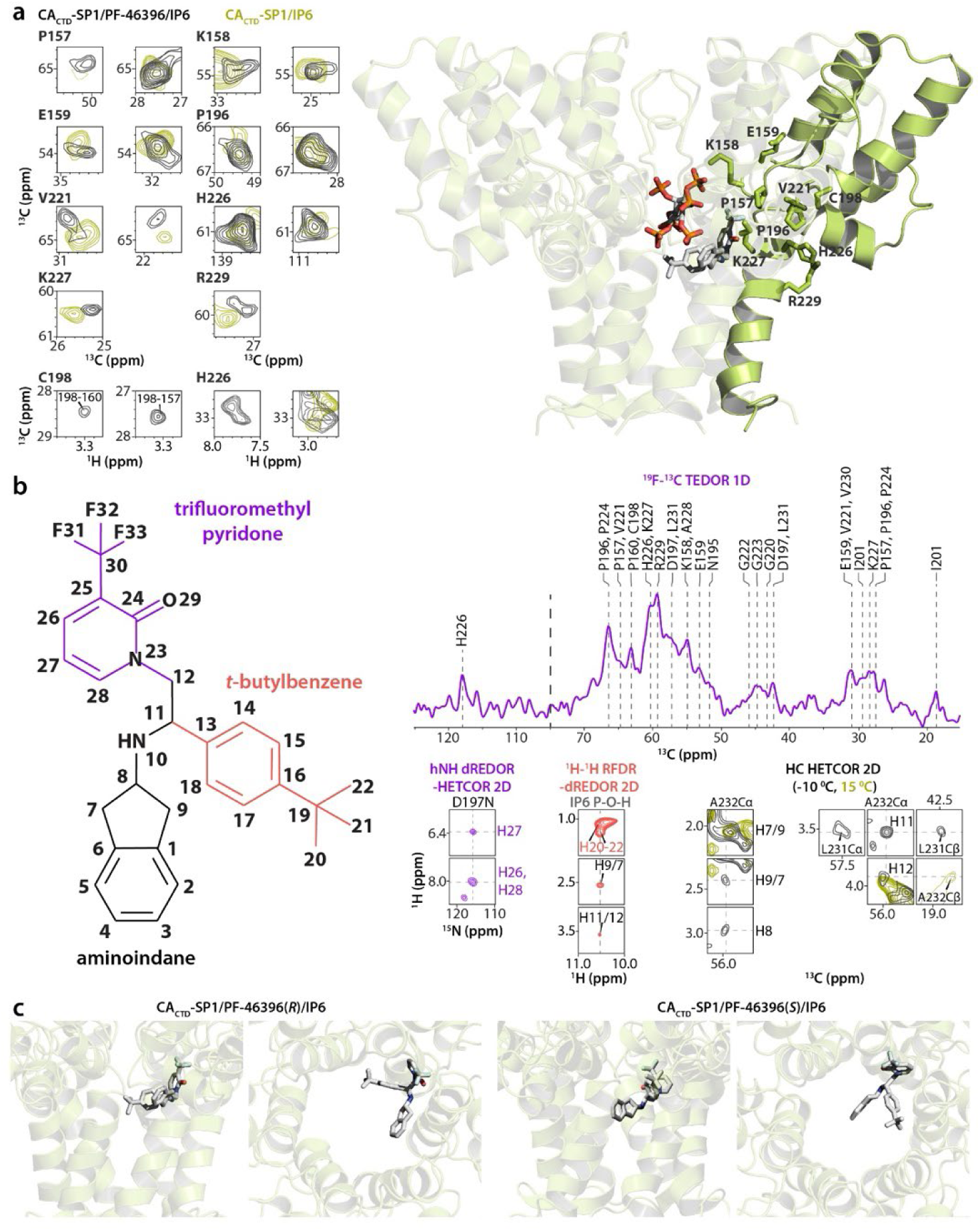
| Distinct binding modes of PF-46396*(S)* and PF-46396*(R)* enantiomers to CA_CTD_-SP1 assemblies. a) Left: superposition of selected regions of 2D CORD 50 ms spectra of U-^13^C,^15^N-CA_CTD_- SP1/IP6 (mustard) and U-^13^C,^15^N-CA_CTD_-SP1/PF-46396(*racem*)/IP6 (gray). MAS frequency was 14 kHz. **Right:** MAS NMR structure of CA_CTD_-SP1/PF-46396(*R*)/IP6; the protein side chains exhibiting chemical shift perturbations upon PF-46396 binding are shown in stick representation. **b) Left:** PF-46396 chemical structure with the aminoindane ring shown in black, the trifluoromethylpyridone unit in violet, and the *t*-butyl- benzene unit in coral. **Right top:** selected regions of 1D ^19^F-^13^C TEDOR spectrum displaying correlations between the PF-46396 trifluoromethyl ^19^F and protein ^13^C resonances. **Right bottom:** selected regions of the 2D hNH dREDOR-HETCOR spectrum illustrating PF-46396 (pyridone ^1^H)-protein (^15^N) correlations; selected regions of the 2D ^1^H-^1^H RFDR-dREDOR spectrum depicting PF-46396 (aminoindane and *t*- butylbenzene ^1^H) and IP6 (P-O-H ^1^H) correlations; superposition of selected regions of the 2D HC HETCOR spectrum of FD-^13^C,^15^N-CA_CTD_-SP1/PF-46396(*racem*)/IP6 displaying PF-46396 (aminoindane ^1^H) and IP6 (^1^H)-protein (^13^C) correlations. The latter spectra were acquired at 15 °C (mustard) and -10 °C (gray). Additional experimental details are summarized in Table S1, Supporting Information. **c) Left:** Top and side views of the binding poses and orientations of PF-46396(*R*) in the 6HB. **Right:** Top and side views of the binding poses and orientations of PF-46396(*S*) in the 6HB.

To unequivocally determine the PF-46396 and IP6 binding mode and orientation in CA_CTD_- SP1 assemblies, we conducted carefully designed correlation experiments. Both PF-46396 and IP6 are at natural ^13^C abundance, making their detection in the NMR spectra challenging due to prohibitively low sensitivity. We, therefore, utilized unique ^19^F and/or ^1^H chemical shifts as NMR reporters to detect the small molecule correlations with isotopically labeled CA_CTD_-SP1. Through a series of 1D and 2D heteronuclear correlation experiments, we were able to unambiguously assign 41 ^19^F-^13^C, ^1^H-^13^C, or ^1^H-^15^N PF-46396-(CA_CTD_-SP1) correlations, which provide distance restraints between the inhibitor and the protein (Fig. 2b). These restraints enabled us to unequivocally determine position and orientation of PF-46396 in the 6HB of the CA_CTD_-SP1 assemblies (Fig. 2c).

In the resulting structures, a PF-46396 molecule occupies a non-centered position within the 6-helix bundle pore, with aminoindane and trifluoromethylpyridone groups facing the protein chains, while the *t*-butylbenzene moiety is oriented toward the 6HB pore. The fluorine atoms of the trifluoromethyl group are in contact with the backbone and side chain carbons of protein residues E159, N195, D197, C198, I201, G220, G223, P224, H226, K227, and R229 (Fig. 2b).

These contacts represent interatomic distances of up to 9.2 Å permitting accurate positioning of the PF-46396 molecule. The unique proton resonances of the pyridone group (PF-46396 H26- 28) exhibit correlations with the backbone nitrogen resonance of D197, and protons H7-9,11,12 of the aminoindane group have correlations to Cα and Cβ resonances of residues L231 and A232. The lack of correlations between resonances of the *t*-butylbenzene moiety and protein side chain resonances agrees with the proposed orientation. Interestingly, correlations between *t*-butyl (PF- 46396 H20-22) resonances, aminoindane (PF-46396 H7/9, H11/12) resonances, trifluoromethyl group ^19^F resonance, and IP6 P-O-H proton resonances, suggest that PF-46396 and IP6 are close to each other, providing direct evidence for simultaneous binding of the cofactor and inhibitor.

### IP6 Adopts Distinct Orientations upon Binding of PF-46396 Enantiomers

In CA_CTD_-SP1/IP6 assemblies, IP6 binds inside the neck region of the channel in the 6HB; its ring is nearly orthogonal to the 6HB axis and parallel to the positively charged side chains of six K158 and six K227 residues, maximizing interactions with those^20^. Remarkably, when bound, IP6 retained some degree of motional freedom with limited rotations inside the 6HB pore on a µs timescale, which, upon BVM binding are arrested and leave IP6 in a tilted orientation^20^. Similarly, PF-46396 binding quenches IP6 dynamics, with the cofactor orientation being enantiomer-dependent, as discussed below.

In the structures of CA_CTD_-SP1 assemblies derived from NMR spectra recorded with different PF-46396 enantiomers, the trifluoromethyl groups occupy identical positions, although the orientations of the aminoindane and *t*-butylbenzene fragments differ. For PF-46396(*R*), the aminoindane and *t*-butylbenzene moieties are nearly parallel to the plane of K227 side chains, while, for PF-46396(*S*), these groups are tilted by about 15 degrees relative to the K227 side chain plane, with the *t*-butyl moiety positioned closer to IP6 and the aminoindane facing SP1 (Fig. 2c). Intriguingly, these small differences in the positions of the PF-46396 enantiomers result in unique IP6 orientations, based on distinct sets of correlations between CA_CTD_-SP1 protons and phosphorus atoms of the cofactor. The IP6–protein contacts were established based on correlations between the IP6 inositol protons and the CA K158 Cε resonance, as well as the IP6 phosphorus and proton resonances of nearby protein side chains, namely, P157, K158, P224, and K227. In total, 20 non-redundant distance restraints were obtained. Finally, five ^1^H-^1^H and one ^1^H-^19^F non-redundant correlations between the P-O-H proton resonance of the cofactor and H11/12, H9/7, H20-22 resonances, as along with the ^19^F resonance of the inhibitor, as well as 14 correlations between IP6 phosphorus resonances and PF-46396(*R*) H2-5, H7/9, H14, H15, H17, and H18, resonances confirmed the simultaneous binding, proximity, and relative orientations of IP6 and PF-46396. These correlations are shown in Figs. 2b and 3 and summarized in Table 1. The distinct P-O-H proton signals observed in multiple spectra indicate the presence of monoanionic IP6 in the samples at pH 8.0, which is often overlooked in the literature^16, 20, 31, 32^ (see Supporting Information for more details). This unique information highlights the power of MAS NMR in providing atomic-resolution structural insights that are currently inaccessible by any other structural biology technique.

**Figure 3.**
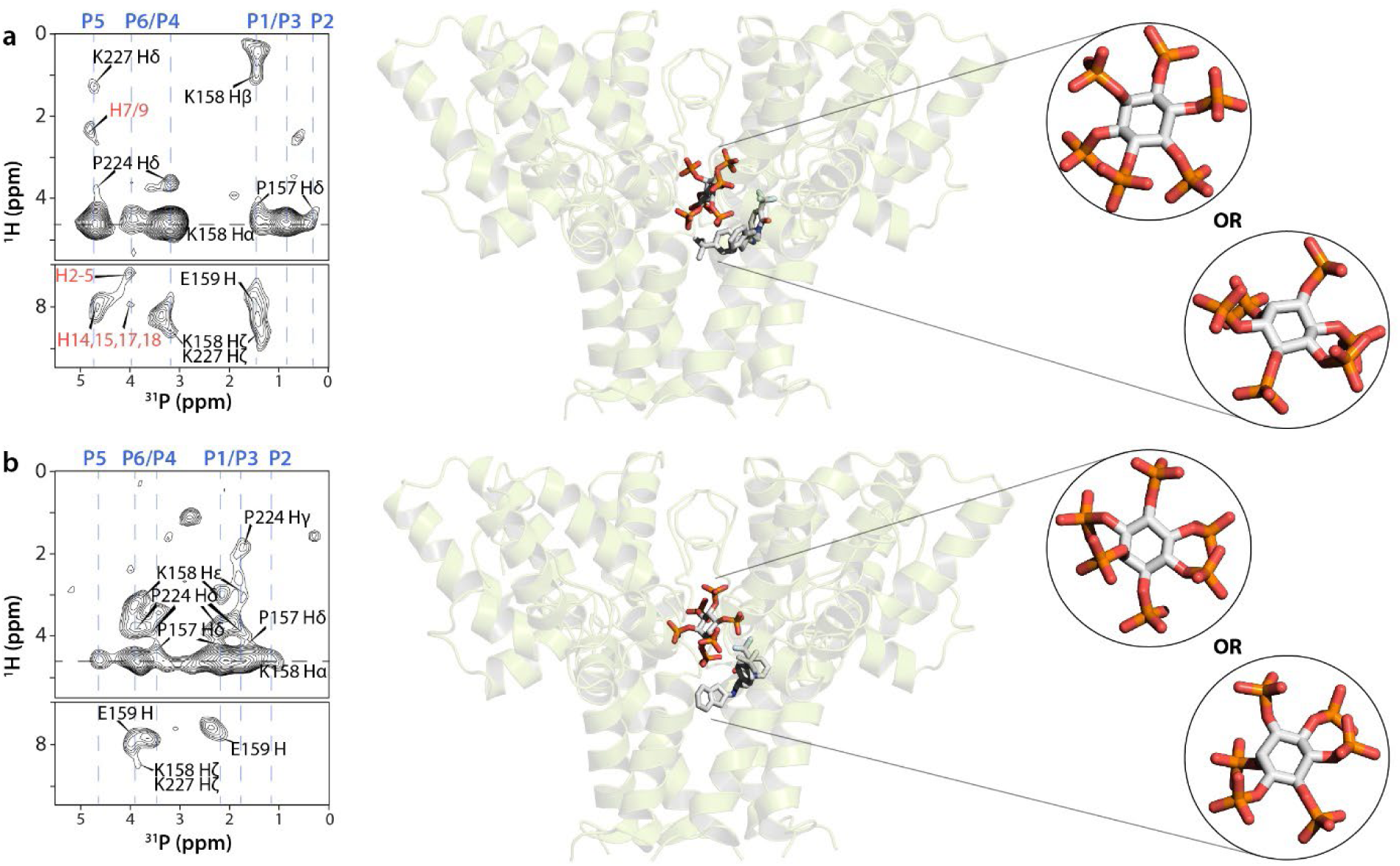
| Distinct orientations of IP6 in the presence of the individual PF-46396*(R)* (a) and PF- 46396*(S)* (b) enantiomers. Left: 2D HP HETCOR spectra of CA_CTD_-SP1/PF-46396(*R*)/IP6 and CA_CTD_- SP1/PF-46396(*S*)/IP6. The assignments of IP6 phosphorus resonances are indicated in blue above each spectrum. The IP6-protein and IP6-PF-46396 contacts are shown in black and coral, respectively. **Middle:** binding poses and orientations of IP6 in the presence of PF-46396(*R*) and PF-46396(*S*). **Right:** expanded views of possible IP6 orientations in the presence of PF-46396(*R*) and PF-46396(*S*).

Upon PF-46396(*S*) binding, IP6 tilts nearly 90 degrees sideways while the P2-P5 C2 symmetry axis remains parallel to the two rings of lysine side chains, permitting the axial P1/6 and P3/4 atoms to make close contacts with the K158 and K227 side chains, respectively (Fig. 3). This orientation is borne out by correlation patterns in the 2D HP HETCOR spectra, where P2 and P5 resonances possess no cross-peaks, except with the K158 Hα resonance, while the resonances of the two pairs of equatorial phosphates (P1/6 and P3/4, located on opposite sides of the C2 symmetry axis) exhibit distinct cross-peak patterns. Surprisingly, upon PF-46396(*R*) binding, the P2-P5 axis of IP6 tilts approximately 60 degrees out of the plane of the lysine side chains, bringing the P5 atom closer to the PF-46396 molecule. This orientation is also supported by the presence of specific correlations between the P5 resonance of IP6 to the aminoindane and benzene proton resonances of PF-46396 (H2-5, H7/9, H14, H15, H17, and H18). All phosphorus atoms possess unique chemical shifts as well as distinct correlation patterns. In addition, the line broadening observed in 2D ^1^H-^31^P HETCOR spectra suggests a coexistence of multiple IP6 orientations, resulting from the different local environments for the phosphorus atoms (Fig. 3). Collectively, all NMR data are consistent with two distinct IP6 orientations in the presence of PF- 46396(*S*) and PF-46396(*R*) enantiomers. For the final structures, each IP6 orientation was refined separately, yielding two distinct bundles of structures for the ternary PF-46396(*S*) and PF- 46396(*R*)/CA_CTD_-SP1/IP6 assemblies.

The unique correlations between CA_CTD_-SP1/IP6 with PF-46396(*S*) and PF-46396(*R*) enantiomers in the ternary assembly permitted the assignment of the absolute configuration of the chiral carbon atom in PF-46396. Two distinct sets of ^1^H-^31^P distance restraints were derived from 2D HETCOR spectra of CA_CTD_-SP1/IP6 assemblies with pure PF-46396 enantiomers, arbitrarily denoted as “set 1” and “set 2” for refinement of the CA_CTD_-SP1/IP6/PF-46396(*S*) and CA_CTD_-SP1/IP6/PF-46396(*R*) structures. Refinement with set 1 resulted in a PF-46396(*S*) orientation consistent with the experimental PF-46396-(CA_CTD_-SP1) cross-peaks. In contrast, refinement with set 2 yielded an orientation that was inconsistent with the experimental data and caused a significant steric clash. In addition, only one of the two ^1^H-^31^P HETCOR spectra exhibits a cross-peak pattern consistent with the IP6 orientation calculated in the presence of PF- 46396(*S*).

For the PF-46396(*R*) enantiomer bound structure, set 2 produced the IP6 orientation without any steric clash between the PF-46396(*R*) and IP6. In contrast, using set 1 resulted in three alternative IP6 binding poses with three too-short distances between PF-46396(*R*) and IP6 on average. Thus, set 2 is only consistent with the PF-46396(*R*) enantiomer/CA_CTD_-SP1/IP6 assemblies.

Overall, each PF-46396 enantiomer/CA_CTD_-SP1/IP6 ternary assembly gives rise to a unique ^1^H-^31^P HETCOR spectrum, enabling us to assign the absolute configuration of the chiral carbon atom.

## DISCUSSION

PF-46396 is a propeller-shaped molecule featuring three distinct chemical blades. Its structure differs significantly from BVM^29^. PF-46396 binds asymmetrically within the 6HB pore of CA_CTD_-SP1/IP6 assemblies, interacting with only three of the six protein chains. Surprisingly, the two PF-46396 enantiomers display distinct binding poses, although similar antiviral activity and infectivity is observed. The absolute configurations of PF-46396(*S*) and PF-46396(*R*) enantiomers were assigned based on their distinct interactions with both CA_CTD_-SP1 residues and IP6. Within the PF-46396(*S*) or PF-46396(*R*)/CA_CTD_-SP1/IP6 ternary assemblies, the IP6 motions are arrested, and the cofactor adopts unique enantiomer-specific orientations. We are not aware of any other instances where a stereospecific assignment of an individual enantiomer of a small molecule in a complex with a protein assembly was determined by MAS NMR.

The position and orientation of IP6 in the presence of PF-46396 explain why PF-46396 rescues infectivity of both K158 and K227 A/I variants that are refractory for IP6 binding, while BVM does not^33^. These BVM escape mutants target the K227 region^6, 21^, while those for PF-46396 cluster in the area surrounding K158^29, 30^. We, therefore, posit that one of the MI escape pathways involves fine-tuning 6HB stability by lowering IP6 binding. As evidenced by our previous CA_CTD_- SP1/IP6/BVM structure^20^, K158 is primarily responsible for coordinating IP6 in the presence of BVM, while our current work shows that IP6 is coordinated by both K158 and K227 in the presence of PF-46396. The preference for PF-46396 escape mutations around K158 (CA G156, P157, and P160)^30^, rather than around the proximal K227 residue, can thus be explained by the distinct PF- 46396 binding position relative to BVM.

## CONCLUSIONS

The atomic details of PF-46396 interactions with CA_CTD_-SP1 and IP6 derived from the current study clarify previous findings on PF-46396 binding and activity. Using PF-46396 structural analogs, the aminoindane and trifluoromethyl moieties were identified as essential groups for binding to Gag, while the *t*-butyl group was shown to be critical for antiviral activity^34^. These findings align with our atomic-resolution structural model, where aminoindane and trifluoromethylpyridone groups interact with protein chains, while the *t*-butylbenzene moiety faces into the 6HB pore.

In addition, our structural model elucidates the mechanism of PF-46396 escape mutations SP1-A1V and SP1-A3V^30^. The CA residues I201, P224, and G225, located near or at the type-II β-turn, as well as H226 in the CA-SP1 junction helix, are all close to the PF-46396 binding site. The SP1-A1 residue is at the edge of the PF-46396 binding pocket, while SP1-A3 resides further down in the helix, and any side chain changes potentially affect PF-46396 binding. Interestingly, it has been established that PF-46396 potency is not reduced in the SP1-V7 BVM-resistance strains, such as SP1-V7A, consistent with PF-46396 binding higher up in the 6HB pore.

Taken together, our results provide a structural explanation for PF-46396 binding, activity, and antiviral escape, opening new strategies for the design of more powerful MIs.

## ACKNOWLEDGEMENTS

This work was supported by the National Institutes of Health (NIH Grant 1U54AI170791 to A.M.G., T.P., and B.K.G-P). Research in the Freed lab is supported by the Intramural Research Program of the Center for Cancer Research, National Cancer Institute, National Institutes of Health. We acknowledge the support of the National Science Foundation (NSF Grant CHE0959496) for the acquisition of the 850 MHz NMR spectrometer. We acknowledge the National Institutes of Health (NIH S10-OD026896A) for the acquisition of the X-ray diffractometer system. We thank Juan R. Perilla and Juan S. Rey for parameterizing the force fields for PF-46396 enantiomers. We are grateful to Ryan W. Russell for the Xplor-NIH structure calculation scripts and Sucharita Sarkar for valuable discussions on structure calculations.

## DATA AVAILABILITY

The MAS NMR atomic structure coordinates have been deposited in the Protein Data Bank under accession codes 9ON8, 9ON9, 9ONA, and 9ONB. The MAS NMR chemical shifts are deposited at the Biological Magnetic Resonance Bank (BMRB) under BMRB entry ID 31245. The crystal structure has been deposited at the Cambridge Structural Database under CCDC 2421271.

## SUPPORTING INFORMATION

2D CORD MAS NMR spectra; summary of MAS NMR samples and experiments; summary of crystal data and structure refinement for PF-46396, viral infectivity assays. This material is available free of charge via the internet at http://pubs.acs.org.

