## Supporting Information for "Structural Basis for HIV-1 Maturation Inhibition by PF-46396 Determined by MAS NMR"

**Table of Contents**

- Supplementary Text: IP6 is present in the monoanionic form in U-^13^C,^15^N-CA_CTD_-SP1/PF-46396/IP6 assemblies.
- Figure S1. 2D CORD 50 ms spectra of CA_CTD_-SP1/IP6 crystalline assemblies with and without PF-46396 bound.
- Figure S2. Superposition of 2D CORD spectra of U-^13^C,^15^N-CA_CTD_-SP1/PF-46396 (*racem*)/IP6 and U-^13^C,^15^N-CA_CTD_-SP1/IP6.
- Figure S3. Correlations for SP1 residues Q6-T12 in 2D CORD spectrum of U-^13^C,^15^N-CA_CTD_-SP1/PF-46396 (*racem*)/IP6.
- Figure S4. Viral infectivity in presence of PF-46396 racemic mixture and pure enantiomers.
- Figure S5. Molecular diagram and labeling scheme of PF-46396 with ellipsoids at 30% probability.
- Figure S6. Representative chain of six H-bonded PF-46396 molecules.
- Figure S7. Unit cell diagram.
- Figure S8. ^1^H MAS NMR spectra of PF-46396 (blue), IP6 (gold), and U-^13^C,^15^N-CA_CTD_-SP1 protein assembly (black).
- Figure S9. Selected regions of ^1^H-^1^H RFDR-dREDOR and ^1^H-^19^F HETCOR 2D spectra.
- Table S1. Summary of MAS NMR experiments.
- Table S2. Crystal data and structure refinement for PF-46396.
- References.
- Author contributions.

**IP6 is present in the monoanionic form in U-^13^C,^15^N-CA_CTD_-SP1/PF-46396/IP6 assemblies**

*Myo-inositol* hexakisphosphate has 12 ionizable phosphate protons (P-O-H) with markedly different dissociation constants. Six protons have pKa values below 2, while the remaining protons' pKa values span a broad range from 5 to 12^1^. During deprotonation, each phosphate group initially loses one proton, converting IP6 from a free acid to a monoanionic form.

Our data support the existence of monoanionic IP6 in the samples at pH 8.0. The ^1^H solid-state MAS NMR spectrum of commercially available IP6 powder (Sigma-Aldrich), acquired without any additional purification, shows two broad signals at 4 and 11 ppm (Figure S8, Supporting Information). The upfield signal is attributed to C-H protons of IP6, while the downfield one corresponds to phosphate (P-O-H) protons. The 2D H-H RFDR-dREDOR and H-F HETCOR spectra (Figure 2b, bottom right panel, and Figure S9, Supporting Information) clearly reveal the presence of PF-46396 H11/12, H9/7, H20-22, and ^19^F resonances aligned with the P-O-H protons of IP6 at 10.5 ppm. No other protons in the samples share this resonance frequency. The closest in chemical shift is W184 H^ε1^ (10.3 ppm^2^), located at the interhexameric interface at least 28 Å from PF-46396 fluorine atoms. Furthermore, the W184 H^ε1^ signal is absent in the dREDOR filtered spectra, as it is directly bonded to a ^15^N-labeled nitrogen in the tryptophan indole ring.

Additionally, the peak pattern for ^31^P signals observed in 2D ^1^H-^31^P HETCOR spectra (Figure 3, left panel) is consistent with that previously described for the monoanionic form of IP6^3^. In that study, Castello et al. identified unique ^31^P resonance patterns characteristic of monoanionic and dianionic IP6. Our spectrum displays the P5 – P4,6 – P1,3 – P2 pattern (from downfield to upfield, marked with blue letters above the spectra) with 1:2:2:1 intensity ratios, which are characteristic of monoanionic IP6. The presence of dianionic IP6 species would produce a P5 – P2 – P1,3 – P4,6 pattern with 1:1:2:2 intensity ratios, which does not match our spectra.

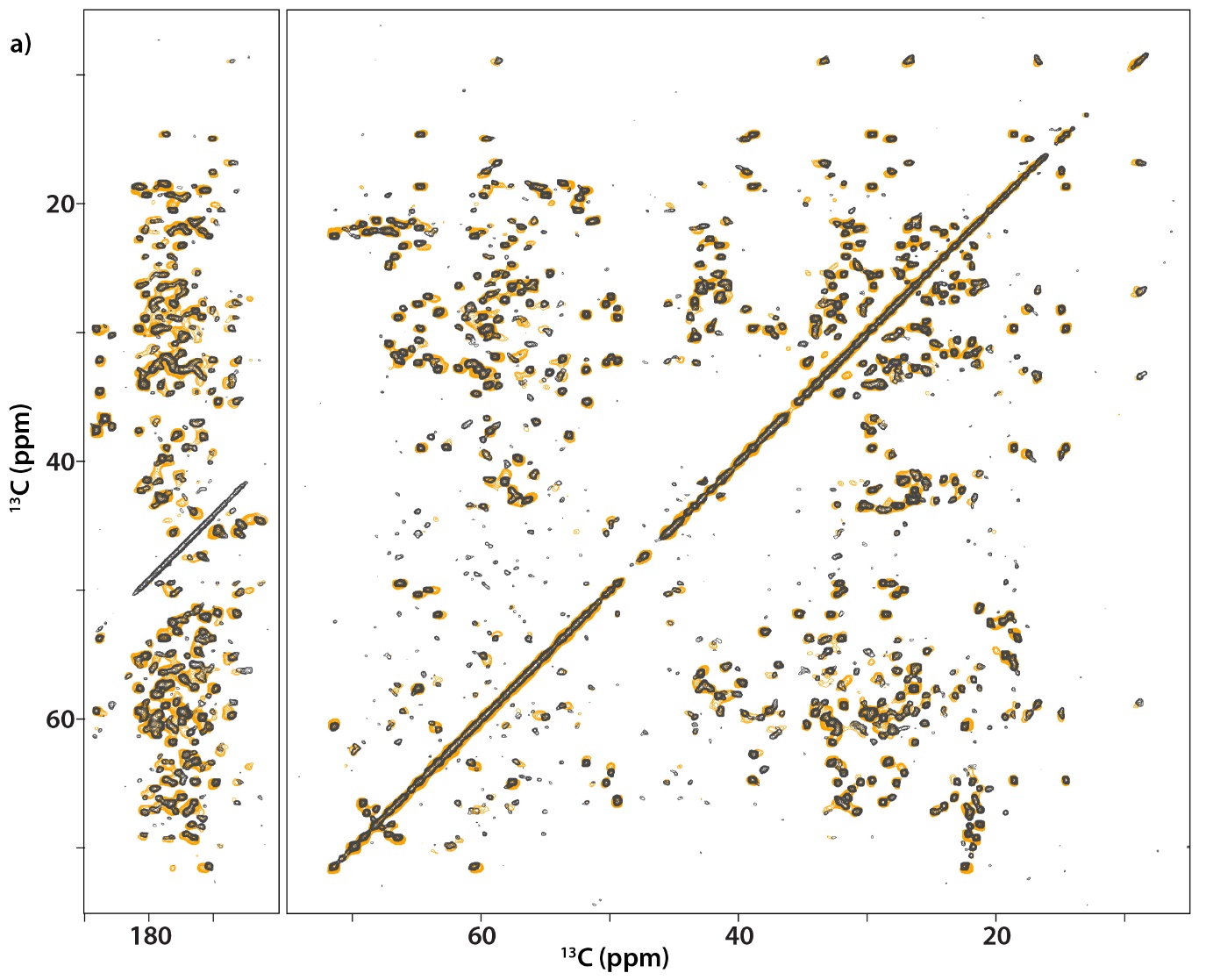

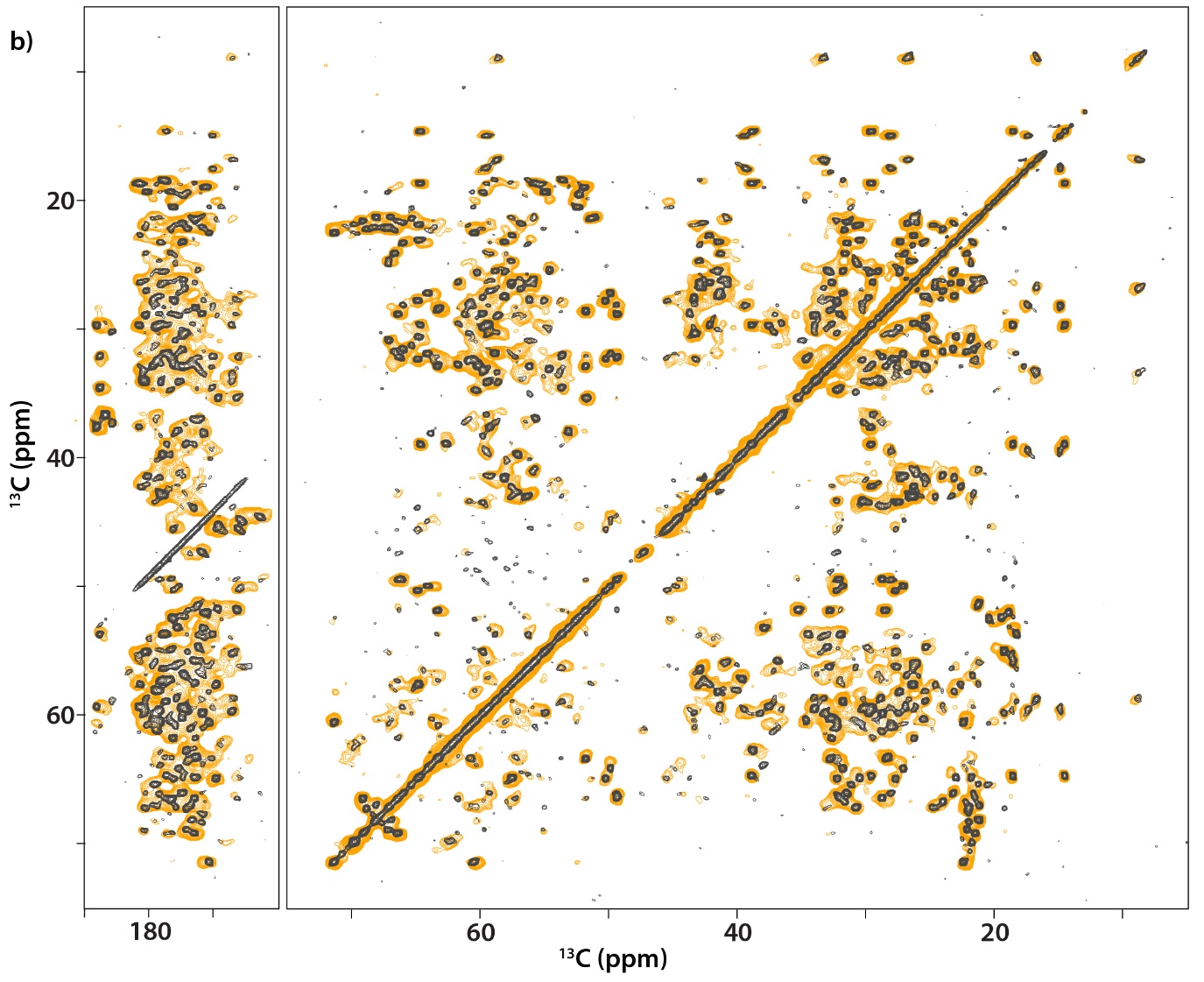

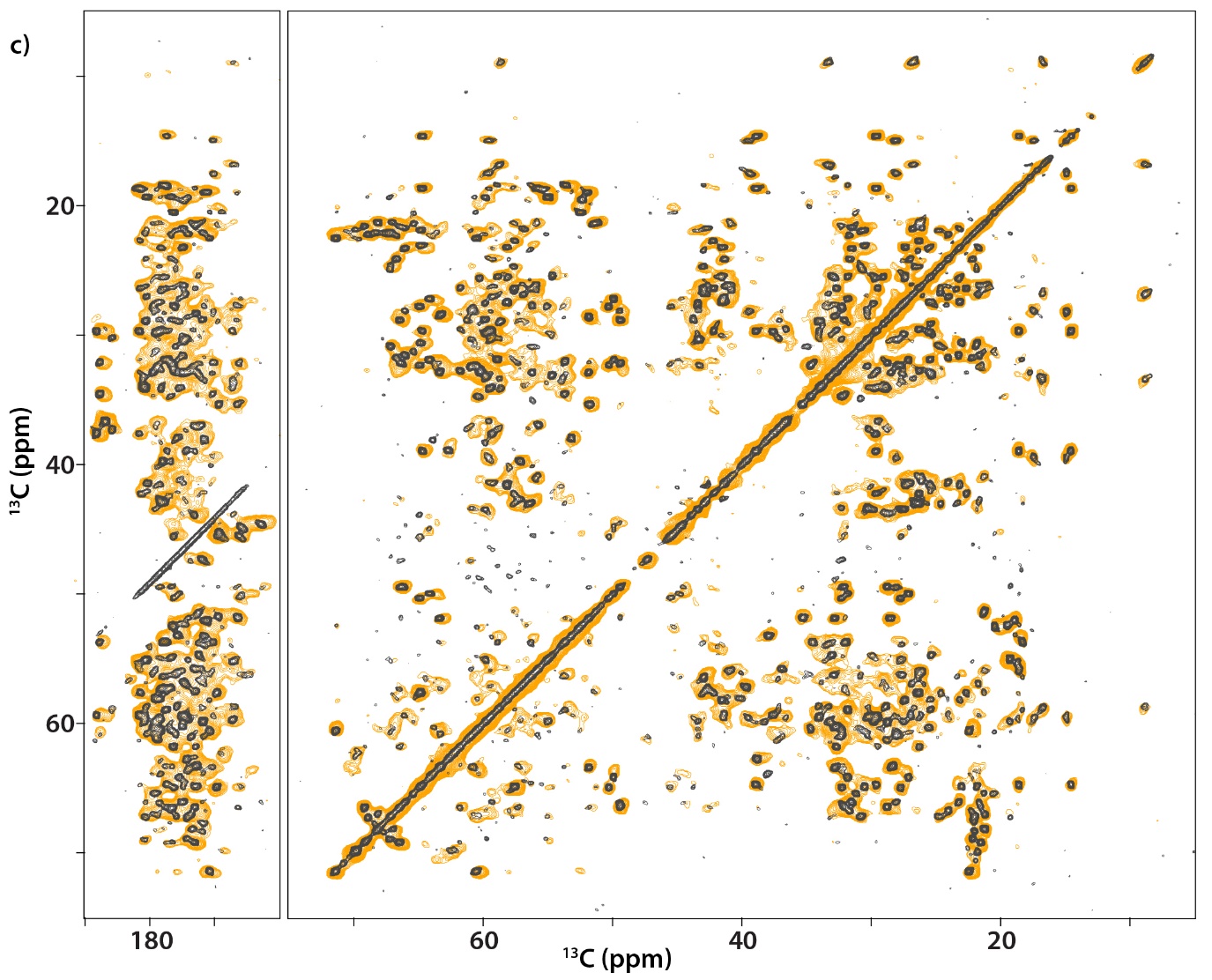

**Figure S1 | 50 ms 2D CORD spectra of CA_CTD_-SP1/IP6 crystalline assemblies with and without bound PF-46396.** Superposition of CORD spectra for U-^13^C,^15^N-CA_CTD_-SP1/IP6 assemblies in the absence (gray) and presence (yellow) of **(a)** PF-46396 (*racem*), **(b)** PF-46396 (*R*), and **(c)** PF-46396 (*S*)/IP6. The spectra were recorded at 14.1 T with a MAS frequency of 14 kHz.

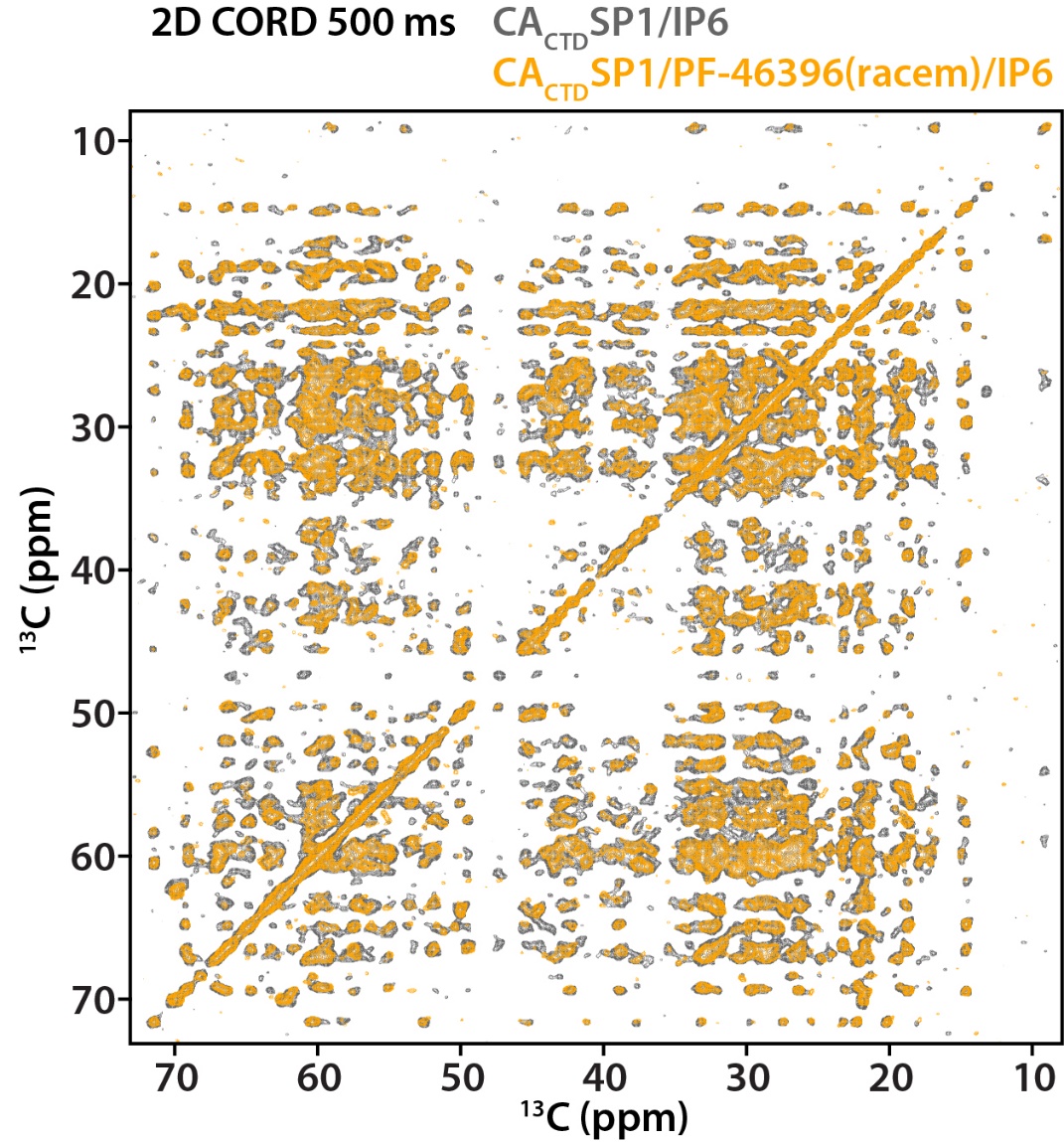

**Figure S2 | Superposition of 2D CORD spectra of U-^13^C,^15^N-CA_CTD_-SP1/PF-46396 (*racem*)/IP6 (yellow) and U-^13^C,^15^N-CA_CTD_-SP1/IP6 (grey).** The spectra were recorded at 14.1 T, with a MAS frequency of 14 kHz. The CORD mixing time was 500 ms.

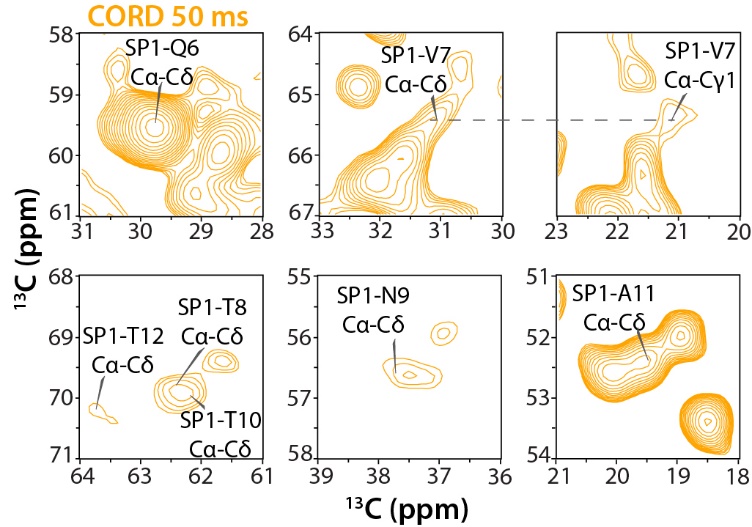

**Figure S3 | Correlations for SP1 residues Q6 toT12 in 2D CORD spectrum of U-^13^C,^15^N-CA_CTD_-SP1/PF-46396 (*racem*)/IP6.** The spectrum was recorded at 14.1 T with MAS frequency of 14 kHz.

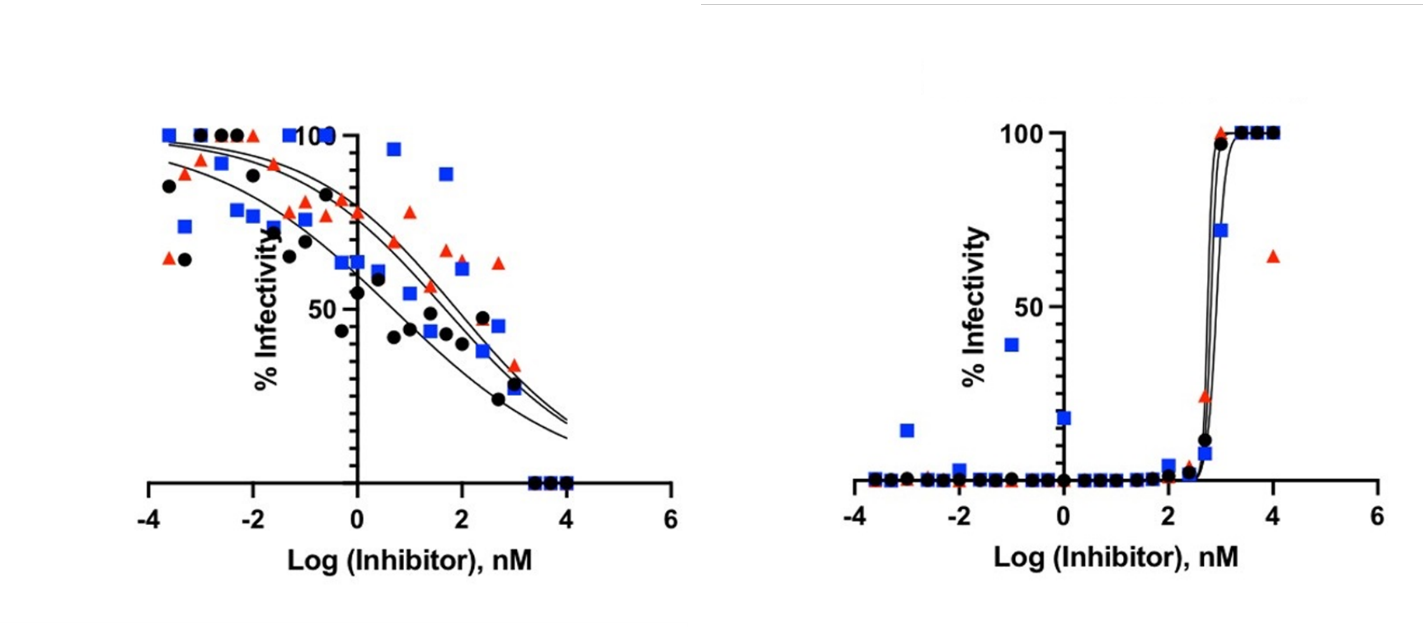

**Figure S4 | Viral infectivity in presence of PF-46396 racemic mixture and pure enantiomers.** Antiviral activity of PF-46396 (*racem*) (black), PF-46396 (*R*) (blue) and PF-46396 (*S*) (red) against HIV-1 WT (left) and the SP1-A3V variant (right).

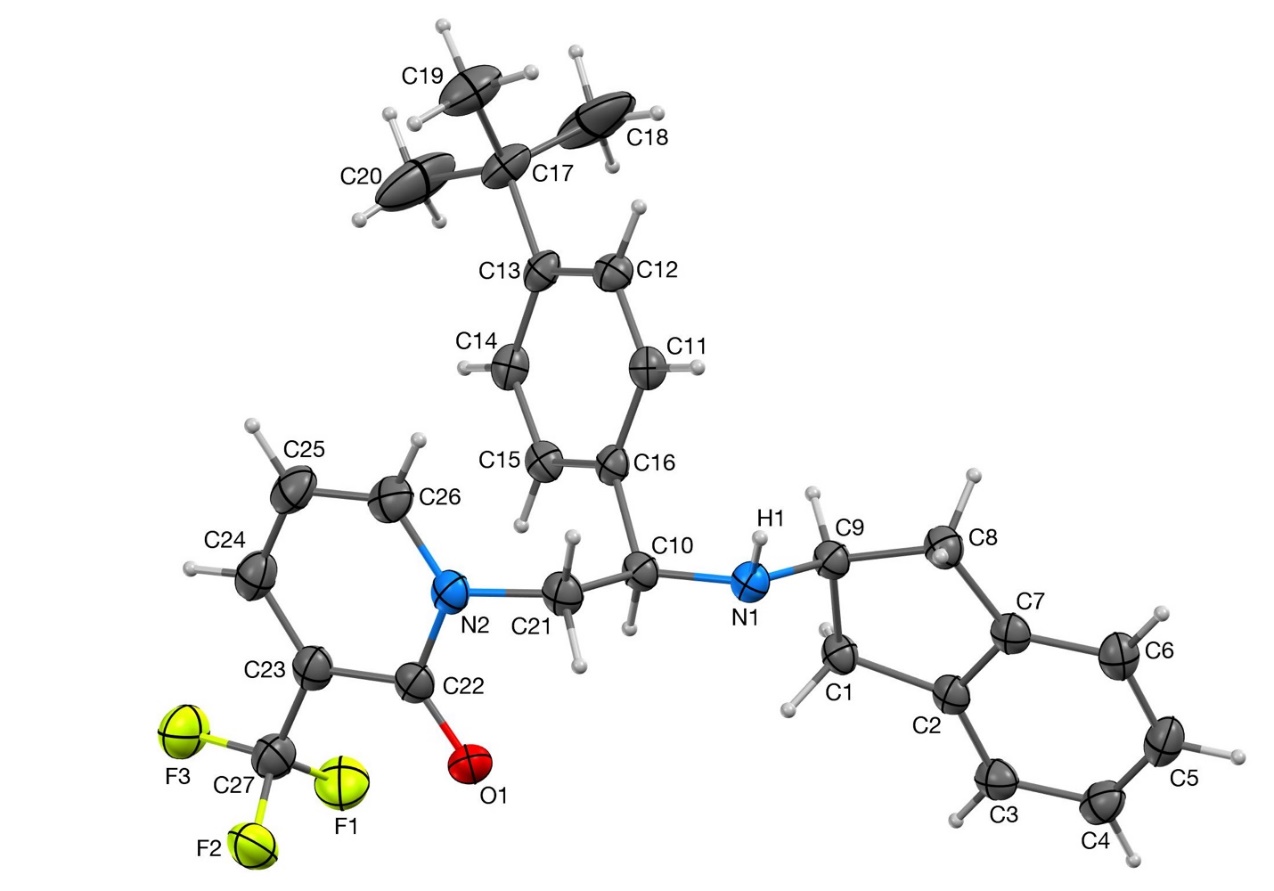

**Figure S5 | Molecular structure of PF-46396 with ellipsoids at 30% probability.** H-atoms depicted with arbitrary radius.

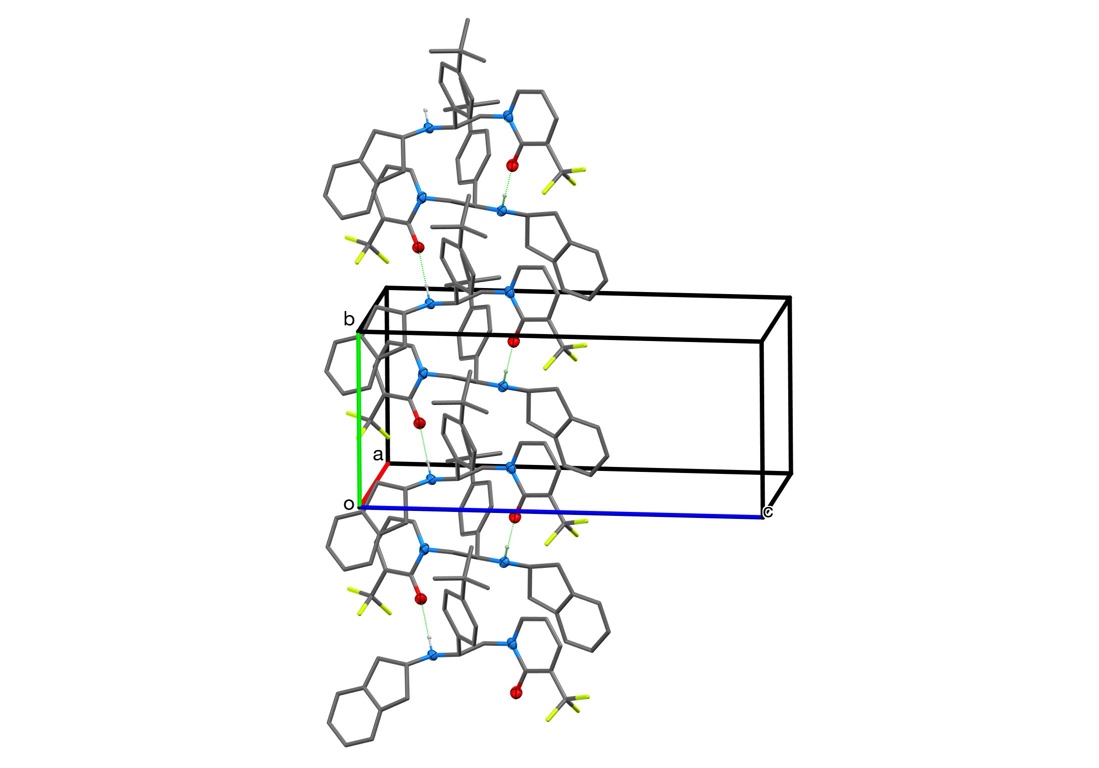

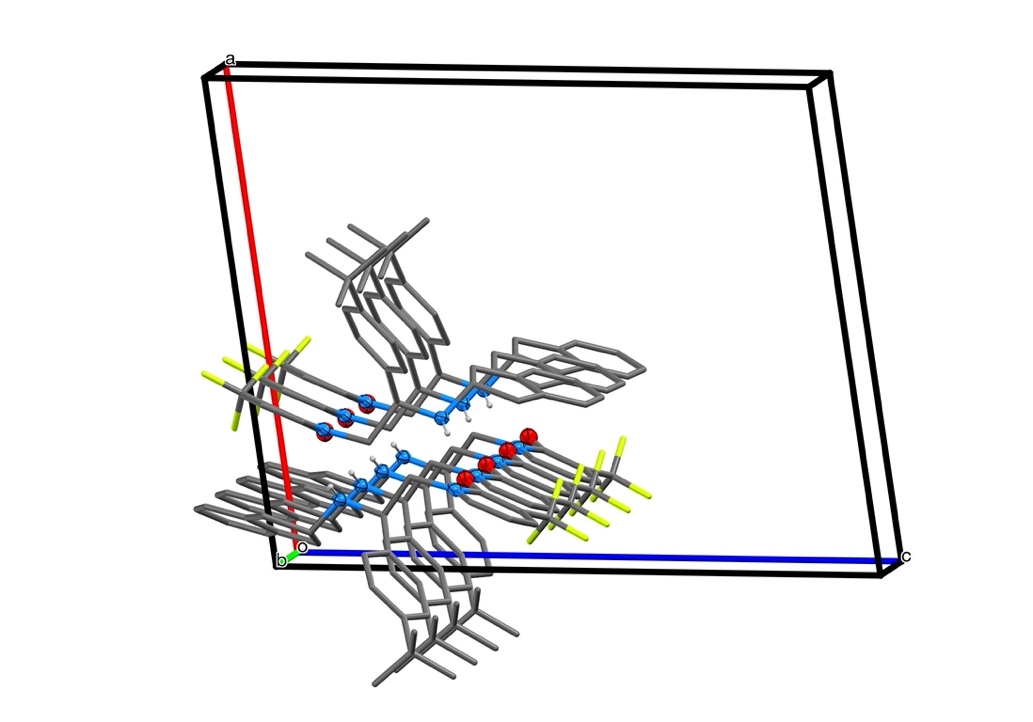

**Figure S6 | Representative chain of six H-bonded PF-46396 molecules** viewed perpendicular to a (top) and b (bottom) showing the 2_1_ screw on b.

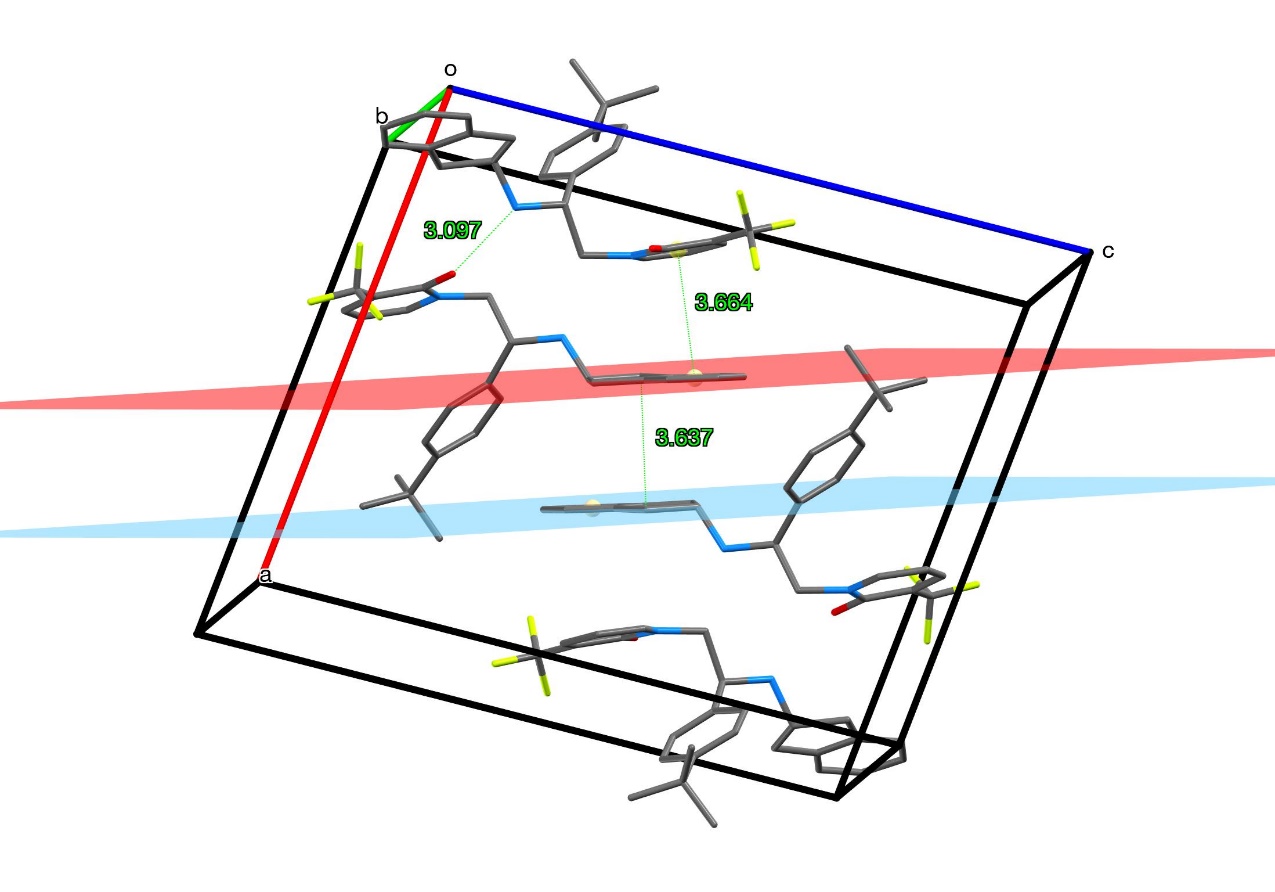

**Figure S7 | Unit cell diagram** of PF-46396 indicating intermolecular distances between the *N*(H)...*O* (3.097 Å), the intrachain pyridone-indane distance (3.664 Å), and the interchain pyridone-pyridone distance (3.637 Å).

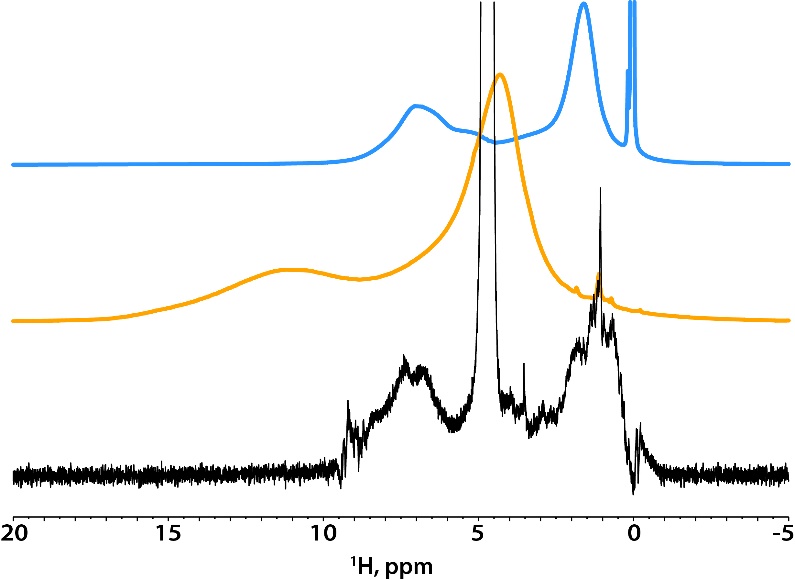

**Figure S8 | ^1^H MAS NMR spectra of PF-46396 (blue), IP6 (gold), and U-^13^C,^15^N-CA_CTD_-SP1 protein assembly (black).** The spectra were recorded at 14.1 T. The MAS frequency was 60 kHz.

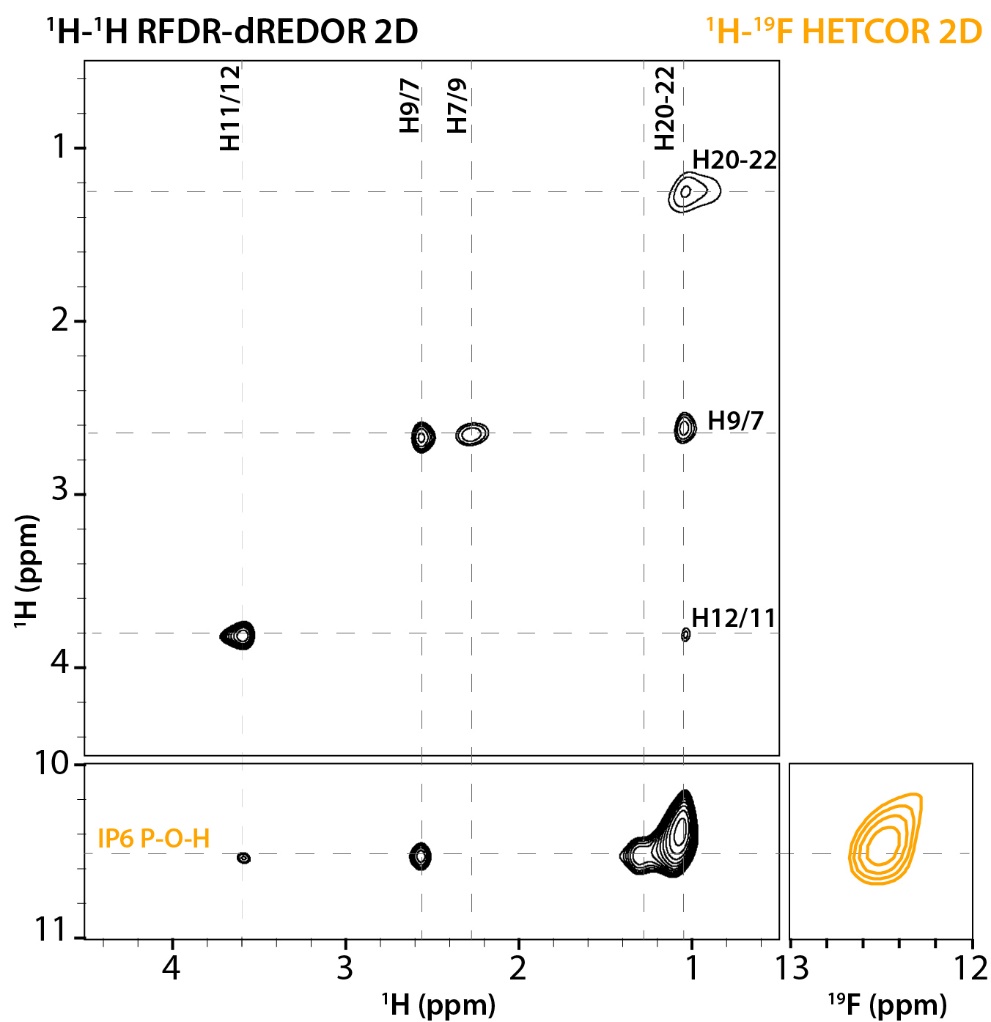

**Figure S9 | Selected regions of ^1^H-^1^H RFDR-dREDOR (black) and ^1^H-^19^F HETCOR (gold) 2D spectra.** Assignments of PF-46396 protons are shown in black, IP6 P-O-H protons – in gold.

**Table S1 |** Summary of MAS NMR experiments.

| **Experimental Parameters** | | | | | | | | | | |
| --- | --- | --- | --- | --- | --- | --- | --- | --- | --- | --- |
| **Sample** | **Experiment** | **B_0_ (T)** | **ω_r_ (kHz)** | **T (±1°C)** | **SNR* (1^st^ FID)** | **NS**** | **t1 points** | **SW1, Hz** | **t2 points** | **SW2, Hz** |
| U-^13^C,^15^N-CA_CTD_-SP1/PF-46396(*racem*)/IP6 | 2D CORD (t_mix_=50 ms) | 14.1 | 14 | 4 | 75 | 96 | 840 | 28000 | 3072 | 45454 |
|  | 2D CORD (t_mix_=50 ms) | 20 | 14 | 4 | 40 | 120 | 840 | 42000 | 2560 | 64102 |
|  | 2D CORD (t_mix_=100 ms) | 14.1 | 14 | 4 | 79 | 96 | 840 | 28000 | 3072 | 45454 |
|  | 2D CORD (t_mix_=250 ms) | 14.1 | 14 | 4 | 55 | 96 | 840 | 28000 | 3072 | 45454 |
|  | 2D CORD (t_mix_=500 ms) | 14.1 | 14 | 4 | 87 | 256 | 840 | 28000 | 3072 | 45454 |
|  | 2D NCACX (t_mix_=50 ms) | 14.1 | 14 | 4 | 28 | 1024 | 130 | 4000 | 3072 | 45454 |
|  | 2D PAIN-CP | 14.1 | 14 | 4 | 26 | 1600 | 104 | 3500 | 3072 | 45454 |
|  | 1D ^19^F-^13^C TEDOR | 11.7 | 60 | 37 |  |  |  |  |  |  |
|  | 2D ^1^H-^19^F TEDOR | 11.7 | 60 | 37 | 23 | 4096 | 30 | 8571 | 4096 | 59524 |
|  | 2D hNH dREDOR-HETCOR | 14.1 | 60 | 33 | 16 | 832 | 40 | 5000 | 2048 | 24038 |
|  | 2D ^1^H-^1^H RFDR-dREDOR | 14.1 | 60 | 33 | 59 | 2048 | 100 | 10000 | 16384 | 24038 |
|  | 1D ^1^H-^31^P CP | 20 | 40 | 15 |  |  |  |  |  |  |
|  | 1D ^31^P Direct | 20 | 40 | 15 |  |  |  |  |  |  |
|  | 2D ^1^H-^31^P HETCOR | 20 | 40 | 15 | 7 | 10240 | 50 | 14000 | 4096 | 138888 |
| FD-^13^C,^15^N-CA_CTD_-SP1/PF-46396(*racem*)/IP6 | 2D ^1^H-^13^C HETCOR | 20 | 14 | -10 | 45 | 256 | 160 | 13333 | 4096 | 64102 |
|  | 2D ^1^H-^13^C HETCOR | 20 | 40 | 15 | 46 | 480 | 160 | 13333 | 4096 | 64102 |
| U-^13^C,^15^N-CA_CTD_-SP1/PF-46396(*R*)/IP6 | 2D CORD (t_mix_=50 ms) | 14.1 | 14 | 4 | 66 | 96 | 840 | 28000 | 3072 | 45454 |
|  | 2D CORD (t_mix_=500 ms) | 14.1 | 14 | 4 | 92 | 384 | 840 | 28000 | 3072 | 45454 |
|  | 2D ^1^H-^31^P HETCOR | 20 | 40 | 15 | 7 | 2048 | 60 | 17000 | 4096 | 138888 |
|  | 1D ^1^H-^31^P CP | 20 | 40 | 15 |  |  |  |  |  |  |
|  | 1D ^31^P Direct | 20 | 40 | 15 |  |  |  |  |  |  |
| U-^13^C,^15^N-CA_CTD_-SP1/PF-46396(*S*)/IP6 | 2D CORD (t_mix_=50 ms) | 14.1 | 14 | 4 | 60 | 288 | 840 | 28000 | 3072 | 45454 |
|  | 2D ^1^H-^31^P HETCOR | 20 | 40 | 15 | 7 | 10240 | 50 | 14000 | 4096 | 138888 |
|  | 1D ^1^H-^31^P CP | 20 | 40 | 15 |  |  |  |  |  |  |
|  | 1D ^31^P Direct | 20 | 40 | 15 |  |  |  |  |  |  |

*SNR is signal-to-noise ratio calculated on the spectrum without apodization.

**NS – number of scans per t1 point

**Table S2 |** Crystal data and structure refinement of PF-46396.

| CCDC | 2421271 |
| --- | --- |
| Empirical formula | C_27_H_29_F_3_N_2_O |
| Formula weight | 454.52 |
| Temperature/K | 100 |
| Crystal system | monoclinic |
| Space group | P2_1_/n |
| a/Å | 15.4606(7) |
| b/Å | 8.1046(4) |
| c/Å | 18.9684(8) |
| α/° | 90 |
| β/° | 98.9870(10) |
| γ/° | 90 |
| Volume/Å^3^ | 2347.60(19) |
| Z | 4 |
| ρ_calc_g/cm^3^ | 1.286 |
| μ/mm^‑1^ | 0.782 |
| F(000) | 960.0 |
| Crystal size/mm^3^ | 0.504 × 0.28 × 0.168 |
| Radiation | CuKα (λ = 1.54178) |
| 2Θ range for data collection/° | 6.872 to 140.142 |
| Index ranges | -18 ≤ h ≤ 18, -9 ≤ k ≤ 9, -22 ≤ l ≤ 23 |
| Reflections collected | 37124 |
| Independent reflections | 4426 [R_int_ = 0.0284, R_sigma_ = 0.0243] |
| Data/restraints/parameters | 4426/0/305 |
| Goodness-of-fit on F^2^ | 1.040 |
| Final R indexes [I>=2σ (I)] | R_1_ = 0.0495, wR_2_ = 0.1277 |
| Final R indexes [all data] | R_1_ = 0.0503, wR_2_ = 0.1283 |
| Largest diff. peak/hole / e Å^-3^ | 0.62/-0.45 |

**AUTHOR CONTRIBUTIONS**

T.P., A.M.G., B.K.G.-P., and O.P. conceived the project and guided the work. R.Z. performed NMR experiments, analyzed the experimental data, and performed the structure calculations. K.K.Z. and B.K.G.-P. prepared samples. C.M.Q. took part in the design and analysis of NMR experiments. S.D.A. and E.O.F. performed infectivity and PF-46396 binding studies. G.P.A.Y. performed X-ray diffraction experiments and determined the X-ray crystal structure. B.J.K., D.S., and C.K. performed PF-46396 chiral separation. R.Z., T.P., and A.M.G. took the lead in writing the manuscript. All authors discussed the results and contributed to manuscript preparation.
